# Amplification of Olfactory Signals by Anoctamin 9 is Essential for Mammalian Olfaction: a Risk Factor for the Covid-19-associated Anosmia

**DOI:** 10.1101/2022.03.02.482745

**Authors:** Hyungsup Kim, Hyesu Kim, Luan Thien Nguyen, Taewoong Ha, Sujin Lim, Kyungmin Kim, Soon Ho Kim, Kyungreem Han, Seung Jae Hyeon, Hoon Ryu, Yong Soo Park, Sang Hyun Kim, In-Beom Kim, Gyu-Sang Hong, Seung Eun Lee, Yunsook Choi, Lawrence B. Cohen, Uhtaek Oh

## Abstract

Sensing smells of foods, prey, or predators determines animal survival. Olfactory sensory neurons in the olfactory epithelium (OE) detect odorants, where cAMP and Ca^2+^ play a significant role in transducing odorant inputs to electrical activity. Here we show Anoctamin 9, a cation channel activated by cAMP/PKA pathway, is expressed in the OE and amplifies olfactory signals. *Ano9*- deficient mice had reduced olfactory behavioral sensitivity, electro-olfactogram signals, and neural activity in the olfactory bulb. In line with the difference in olfaction between birds and other vertebrates, chick ANO9 failed to respond to odorants, whereas chick CNGA2, a major transduction channel, showed greater responses to cAMP. Importantly, single-cell transcriptome data from Covid-19 patients revealed that *Ano9* transcripts were markedly suppressed among genes in the olfactory signal pathway. The signal amplification by ANO9 is essential for mammalian olfactory transduction, whose downregulation may be a risk factor for the olfactory dysfunction in Covid-19 patients.

## (Introduction)

Olfaction is critical for the survival of vertebrates. If the animals do not discriminate against the smells of prey or predators, they cannot survive. Similarly, because olfactory cues are important in suckling, the pups will not survive or grow poorly if they are anosmic ^1^. The olfactory epithelium (OE) in the nasal cavity is a specialized epithelium where odorants bind odorant receptors in olfactory sensory neurons (OSNs) for sensory transduction. OSNs are bipolar neurons that transduce odorants to electrical signals and convey the electrical signals to olfactory bulbs (OB) ^2^.

Earlier studies reported that the cAMP signaling pathway plays a crucial role in detecting odorants ^3^. Odorant binding to receptors in the cilia of OSNs stimulates adenylyl cyclase to synthesize cAMP, which opens CNG channels and thereby leads to the depolarization of OSNs ^4^. Disruption of the adenylyl cyclase type III or *Cnga2*, a major subunit of the CNG channel complex, leads to anosmia and reduces the odorant-evoked electro-olfactogram (EOG) response ^5, 6^.

Odorants or cAMP increase intracellular Ca^2+^ in the OE ^7, 8^, which opens Ca^2+^- activated Cl^-^ channels for prolonged depolarization ^9–11^. The source of Ca^2+^ is thought to be mediated by CNG channels. However, intracellular Ca^2+^ desensitizes the CNG channels ^12, 13^. Thus, a question remains on how intracellular Ca^2+^ can be maintained to open the Ca^2+^-activated Cl^-^ channel. Therefore, a source of cAMP-evoked intracellular Ca^2+^ other than CNG channels may be necessary.

In the present study, we identified another cAMP-activated cation channel, Anoctamin 9 (ANO9/TMEM16J), that is highly expressed in the OE and amplifies CNG channel-initiated electrical signals in OSNs. The signal amplification by ANO9 is obligatory as animals become less sensitive to odors when *Ano9* is ablated. Also, we found that birds may not need the amplification by ANO9 as chick ANO9 is non-functional but chick CNG channels are more sensitive to cAMP, suggesting the diversity in olfactory signals among species.

Because one of the major symptoms of the Covid-19 is hyposmia or anosmia ^14–17^, we also determined if changes in ANO9 expression contributes to the pathology.

## (RESULTS)

### The genetic ablation of *Ano9* impairs odor discrimination

We firstly sought to see if ANO9 is expressed in the OE. Dense ANO9 immunoreactivity was observed in the OE of the wild-type (WT) but not in the *Ano9* ablated mice, specifically at the cilia of OE, where it co-localized with an olfactory marker protein (**Fig. 1a, b**). The ANO9 localization at the ciliary region, an apical surface of the OE where olfactory signals begin with olfactory receptors, was further confirmed by its co-localization with a neuronal marker, Tuj-1 and a cilia-specific marker, acetylated tubulin ^18–20^ (**Fig. 1c**). The presence of *Ano9* mRNA along with transcripts of main olfactory signaling genes and Anoctamins in the mouse and human OE was quantified by a quantitative PCR (**Extended Data Fig. 1**). *Ano9* transcripts were observed in both human and mouse OE. Its expression levels were relatively constant between the two species. However, the expression patterns of other olfactory signal genes such as *Cnga2, Cnga4, Omp*, and *Ano2* were different between the two species (**Extended Data Fig. 1**).

**Fig. 1:**
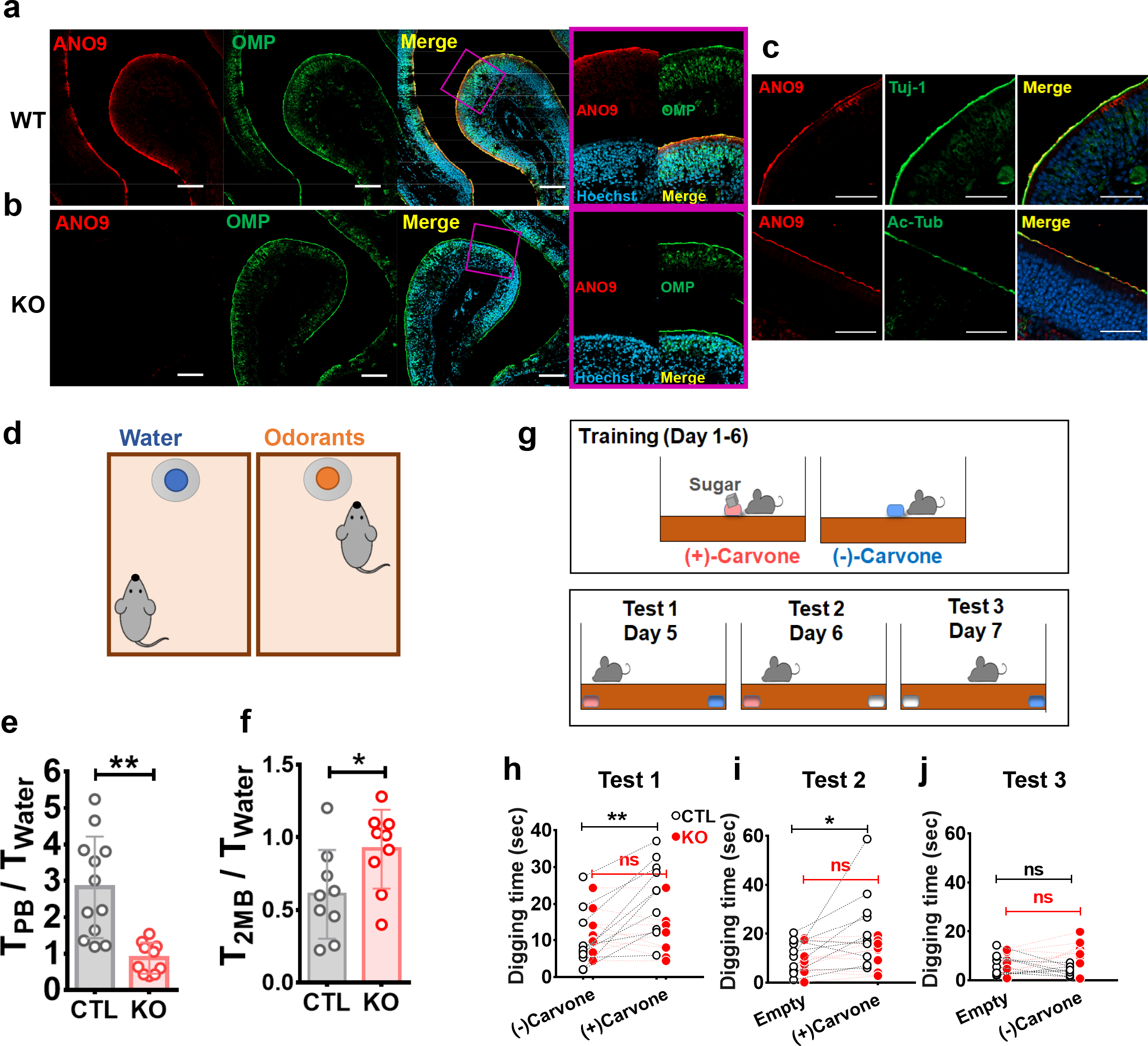
ANO9 expressed in the olfactory epithelium is important for odor recognition. **(a),** Immunofluorescence images of ANO9 in the mouse OE. Sections of the nose of WT and KO **(b)** mice were stained with anti-ANO9 or olfactory marker protein (OMP) antibody **(a-b)**; images of WT epithelia stained with Tuj-1 and acetylated tubulin (Ac-Tub) antibodies are shown in **(c)**. Hoechst 33342 was used for nuclear staining. The right panels in panel **a** and **b** show magnified images of the square areas in the third panel. Scale bars: 100 μm. **d-f,** Scheme of odorant preference test **(d)**. Exploring time was measured in a targeting zone near a dish containing peanut butter (PB), 2-methylbutyric acid (2MB), or water. The ratio of time spent exploring the dish with PB **(e)** or 2MB **(f)** versus water. CTL: control littermates. * p < 0.05, ** p < 0.01, Student T-test **g,** Scheme of odor discrimination learning test (see **Methods**) **h-j,** Summaries of the time taken by mice to dig at each end of the test cage, where (+) carvone and/or (-)-carvone were buried. Note that the CTL littermates spent a longer time digging the place where (+)-carvone was buried. ns: not significant. * p < 0.05, ** p < 0.01, Two-way ANOVA followed by Bonferroni test.

To determine the role of ANO9 in olfaction, we constructed *Ano9* floxed (*Ano9*^fl/fl^) mice that were crossed with CMV-cre transgenic mice to delete *Ano9* in the whole body, including the OE. Pups of CMV-cre;*Ano9^fl/fl^* (knock-out, KO) mice appeared healthy during the neonatal period. However, *Ano9* KO mice appeared hyposmic to odor-preference tests. Firstly, we measured the time spent at places where peanut butter, 2-methylbutyric acid, or water was set in a dish with a perforated lid ^21^ (**Fig. 1d**). Peanut butter is known to be attractive to mice and 2-methylbutyric acid repulsive to mice ^21^. Littermate control (*Ano9^fl/fl^*: CTL) mice spent more time exploring the dish where peanut butter was placed than the dish where water was placed. However, *Ano9* KO mice failed to show a preference for peanut butter (**Fig. 1e**). Similarly, CTL mice showed an aversive reaction to 2-methylbutyric acid as they spent a shorter time exploring the 2- methylbutyric acid-placed location than the water-placed location. However, *Ano9* KO mice failed to avoid the aversive smell as the search time for 2-methylbutyric acid was almost equal to that for water (**Fig. 1f, Extended Data Fig. 2b**).

**Fig. 2.**
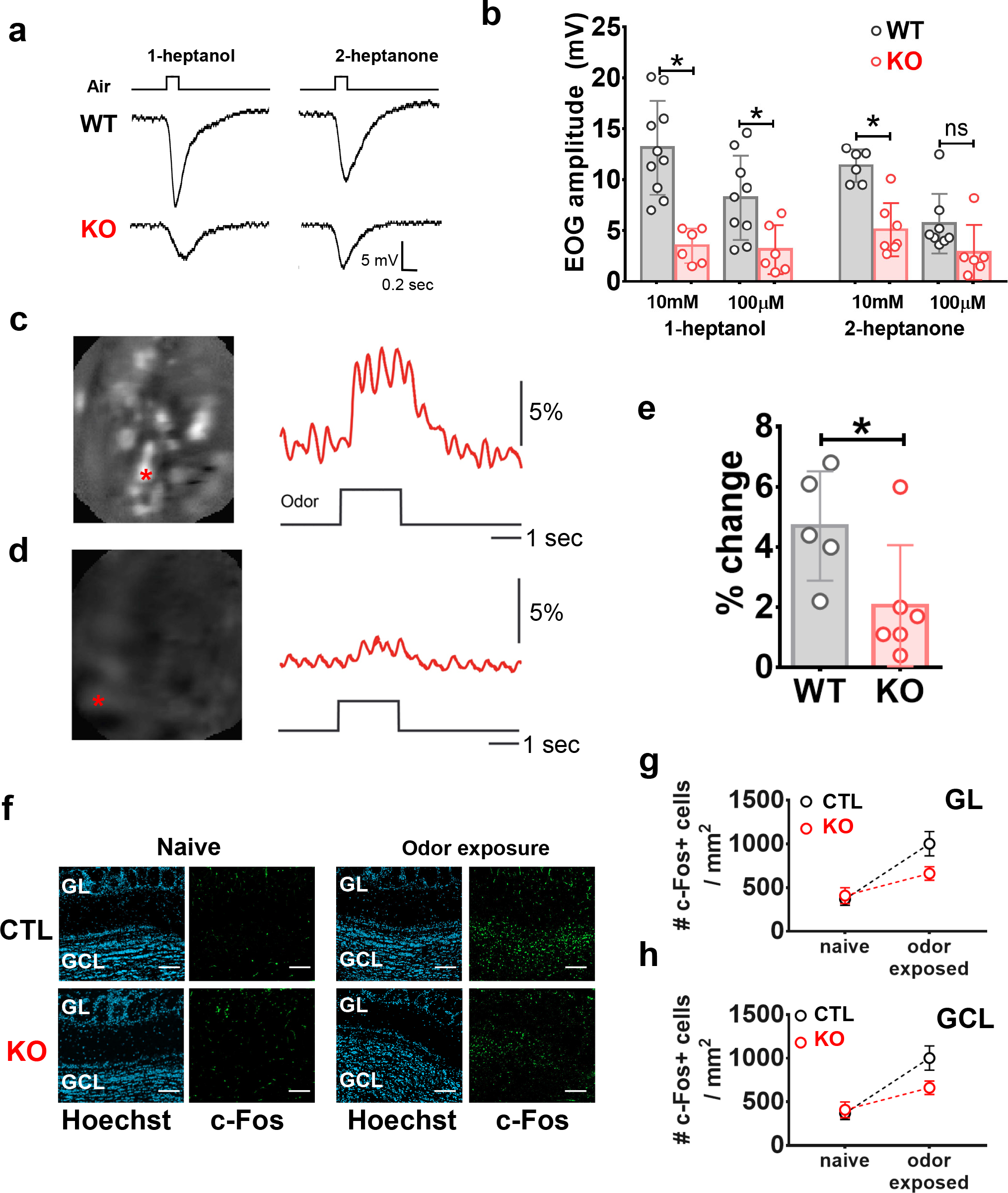
Genetic deletion of *Ano9* reduces EOG responses and in-vivo OB responses to odorants. **a,** Representative electro-olfactogram (EOG) traces of WT and KO mice in response to air pulses containing 10 mM 1-heptanol or 2-heptanone. The field potentials of OSNs in response to odorant-containing air puffs were measured at the olfactory turbinates. **b,** Summary of EOG signals of WT and *Ano9* KO mice in response to 0.1 and 10 mM 1-heptanol or 2-heptanone. ns: not significant, * p < 0.05, Student T-test **c-d,** Representative images of the Ca^2+^ increase in olfactory bulb glomeruli of WT **(c)** and *Ano9* KO **(d)** mice in response to an odor presentation (right panel). The olfactory bulbs were loaded with a Ca^2+^-sensitive dye, Oregon Green 488 BAPTA-1 AM. Odorants were presented to the nose for 2 sec. Right panel: The time courses of the Ca^2+^ response to the odor presentation in two of the olfactory bulb glomeruli (indicated by the red asterisks in the left panel). The traces were low-pass filtered with a 3 Hz Gaussian filter. The scale bar on the right indicates the size of a 5% increase in fluorescence. **e,** Summary of the largest fractional fluorescence change in the olfactory bulb of each mouse in response to odorants. Because most of the Ca^2+^ signals in three out of five KO mice were smaller than the background signals, we used only the largest fluorescence change in each mouse for the statistical analysis. * p < 0.05, Student T-test. **f,** c-Fos expressions in the olfactory bulb of CTL and *Ano9* KO mice after 30-min exposure to odorants. GL; glomerular layer, GCL; granular cell layer, Scale bars: 100 μm. **g-h,** Summary of the number of c-Fos positive cells in the GL **(g)** and GCL **(h)** in the olfactory bulb before and after 30-min exposure to odorants. GL; p < 0.001, F(1, 18) = 20.1, GCL; p < 0.01, F(1, 18) = 13.3, Two-way ANOVA.

Next, we further investigated whether ANO9 would be required for odor discrimination ^22, 23^. Two carvone enantiomers were used to determine if mice could discriminate between (+)-carvone from (-)-carvone (**Fig. 1g**). Littermate CTL and *Ano9* KO mice were trained for four days to learn that (+)-carvone is associated with a sugar reward, whereas sugar is not rewarded for mice exposed to (-)-carvone. On day 5 (Test 1), the two isomers were placed separately at each end of the cage under the bedding. At this time, sugar was not rewarded for choosing (+)-carvone. The time spent digging at each place was measured. CTL mice spent significantly more time digging the bedding where (+)-carvone was placed than digging the bedding buried with (-)-carvone (**Fig. 1h**). In contrast, *Ano9* KO mice spent equal time digging in each place (**Fig. 1h**). On day 6 (Test 2) and day 7 (Test 3), an enantiomer and empty dishes were buried. In these tests, CTL mice spent a longer time digging the place where (+)-carvone was buried than the empty place, whereas *Ano9* KO mice did not (**Fig. 1i**). In addition, both genotypes failed to show a preference for digging at the place with (-)-carvone (**Fig. 1j**). These results indicate that the genetic deletion of *Ano9* impairs odor discrimination.

### *Ano9* KO mice show a reduced EOG response and reduced *in-vivo* neural activity of the OB to odorants

To determine the odor-evoked activity of OSNs of wild-type (WT, C57BL/6) and *Ano9* KO mice, the electro-olfactogram (EOG), the field potentials of OSNs in response to odorant-containing air puffs were measured ^3, 6^. For olfactory stimuli, common volatile odorants, 1-heptanol and 2-heptanone, were used ^24^. EOG voltage responses to odorant-containing air puffs were recorded from olfactory turbinates. Puffs of the air containing 1- heptanol and 2-heptanone evoked robust field potentials in the olfactory turbinates of WT mice in a dose-dependent manner (**Fig. 2a, b**). In contrast, the EOG responses of the olfactory turbinates isolated from *Ano9* KO mice were markedly reduced (**Fig. 2a, b**).

We then compared the *in-vivo* olfactory responses to odorants of WT and *Ano9* KO mice by measuring calcium signals in the olfactory bulb. As the glomeruli in the olfactory bulb receive innervation from OSNs ^25^, we measured odorant responses in WT and *Ano9* KO mice by imaging Ca^2+^ signals in the olfactory bulb glomeruli that were loaded with the Ca^2+^-sensitive dye, Oregon Green 488 BAPTA-1, AM. Odorant stimulation to the nose elicited Ca^2+^ signals in the glomeruli of WT mice (**Fig. 2c**).

However, the glomeruli activity map from an *Ano9* KO mouse (**Fig. 2d**) was considerably dimmer than those from a WT mouse (**Fig. 2c**). The amplitudes of the odor-evoked calcium signals in *Ano9* KO mice were significantly smaller than those of the WT mice (**Fig. 2e**).

c-Fos expression is a useful method for mapping neuronal activities in various brain regions because c-*fos*, an immediate early gene, is expressed relatively rapidly in active neurons ^26^. Therefore, the immunofluorescence of c-Fos was assayed to determine the activity of olfactory bulb neurons stimulated by odorants ^27, 28^. Induction of c-Fos in the olfactory bulb was observed in CTL mice after 30 min exposure to odorant mixtures (**Fig. 2f**). In CTL mice, the nasal odorant mixture induced a 2.7- and 4.9-fold increase in c-Fos expression in the glomerular layer and the granular cell layer of the olfactory bulb, respectively (**Fig. 2g, h**). However, in *Ano9* KO mice, the odorant stimulation increased c-Fos expression in glomerular and granular cell layers but significantly less than those of CTL mice (**Fig. 2g, h**).

### ANO9 amplifies the cAMP-evoked CNG channel currents

We determined whether ANO9 contributes to cAMP-evoked currents in freshly-isolated OSNs from the mouse OE. Whole-cell currents were recorded from isolated OSNs with a pipette containing 100 μM cAMP. Ca^2+^-activated chloride channel blockers, 4,4’-diisothiocyanatostilbene-2,2’-disulfonic acid (DIDS) and niflumic acid, were added to the bath solution to block the anion currents through ANO2 in the OSNs ^10, 11, 29^. As shown in **Fig. 3a**, intracellular cAMP-induced rapid and robust currents with an average amplitude of 962 ± 130 pA (n = 9) at -50 mV in OSNs isolated from WT mice. In contrast, OSNs isolated from *Ano9* KO mice elicited the cAMP-evoked currents with markedly reduced amplitudes (270 ± 45 pA, n = 9) (**Fig. 3a, b**).

**Fig. 3.**
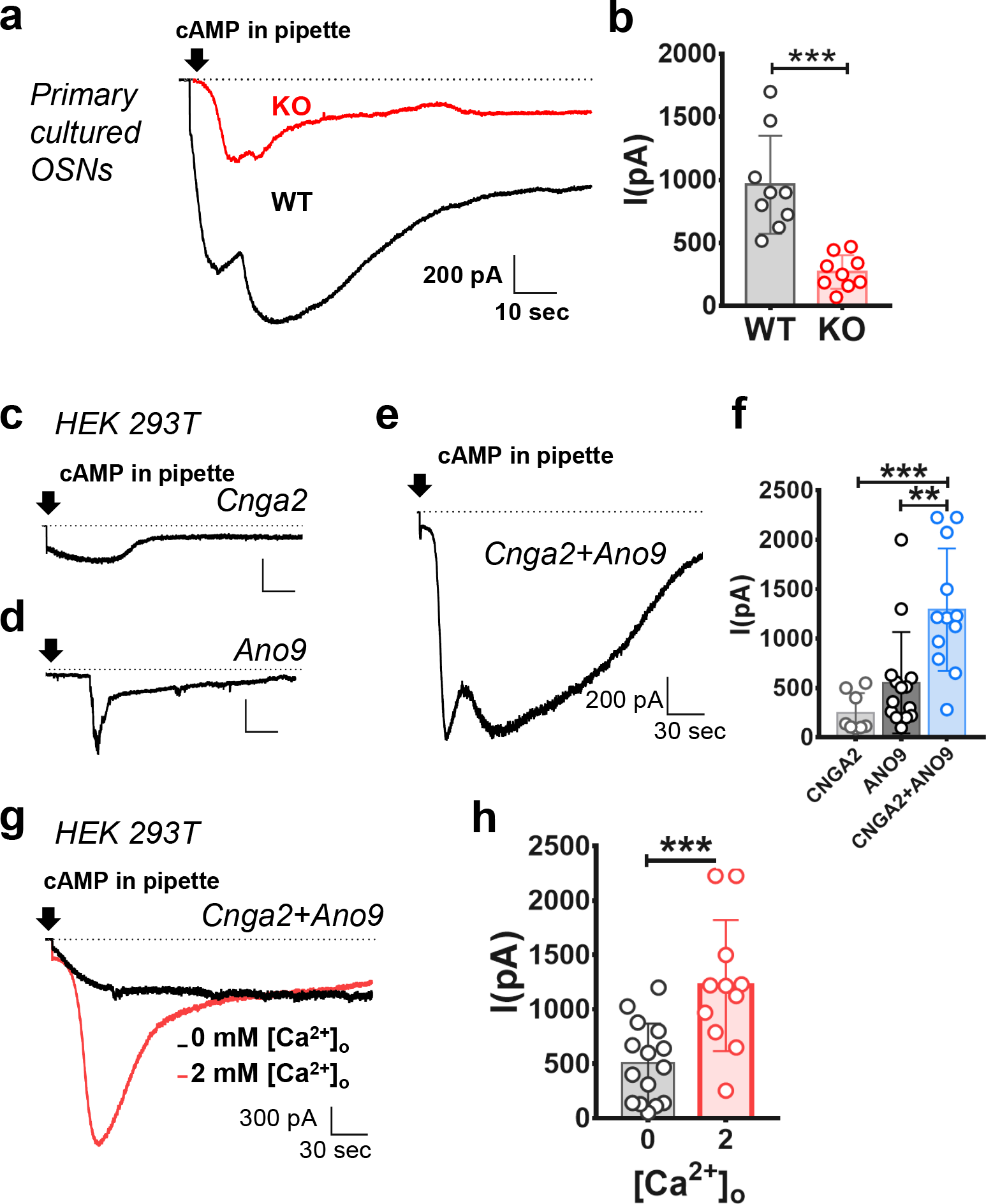
ANO9 amplify CNG currents in OSNs. **a,** The cAMP-induced whole-cell currents of primary cultured OSNs isolated from WT and *Ano9* KO mice. The pipette solution contained 100 μM cAMP. Ehold = -50 mV. The arrow indicates the time that the whole-cell condition was obtained. **b,** Summary of the amplitude of the cAMP-induced whole-cell currents of OSNs from WT and *Ano9* KO mice. *** p < 0.001, Student T-test **c-f,** The cAMP-induced currents of HEK 293T cells transfected with *Cnga2* **(c)**, *Ano9* **(d)**, or *Cnga2*+*Ano9* **(e)**. Intracellular Ca^2+^ was kept at 1 μM. **(f)** Summary of the cAMP-induced current amplitudes. ** p < 0.01, *** p < 0.001, One-way ANOVA, Tukey’s post-hoc test. **g-h,** Representative traces **(g)** and summary of the amplitudes **(h)** of the cAMP-induced currents of HEK 293T cells transfected with *Cnga2+Ano9* recorded in 0 or 2 mM Ca^2+^ in the bath solution. *** p < 0.001, Student T-test.

We then tested whether ANO9 can amplify the cAMP-induced CNG channel currents in a heterologous system. The CNGA2 subunit, one of the main subunits of olfactory CNG channel complex ^30, 31^, can form functional ion channels on its own when expressed heterologously ^32^. In HEK 293T cells transfected with *Cnga2*, 100 μM cAMP in the pipette evoked relatively small currents (**Fig. 3c, f**). The cAMP also evoked currents in ANO9/HEK cells but with a significant delay (58 ± 22 sec, n = 15). However, the cAMP evoked much larger currents without delay in HEK 293T cells transfected with *Cnga2* and *Ano9* (**Fig. 3e, f**). The augmenting capacity of ANO9 was Ca^2+^-dependent as the cAMP-evoked currents in HEK 293T cells transfected with *Cnga2* and *Ano9* were markedly reduced when the external Ca^2+^ was removed (**Fig. 3g, h**).

### ANO9 is a cation channel activated by the cAMP/PKA pathway

As previously reported ^33^, ANO9 was activated by intracellular cAMP (100 μM). But, the cAMP-evoked currents were blocked by a PKA inhibitor, H-89, suggesting that ANO9 is gated by phosphorylation by PKA ^33^. To confirm its activation by PKA, we applied cAMP and a purified catalytic subunit of PKA to inside-out membrane patches of HEK 293T cells transfected with *Ano9* (ANO9/HEK). As shown in **Fig. 4a**, the application of 100 μM cAMP alone to an inside-out membrane patch of ANO9/HEK cells failed to activate single-channel currents. However, when purified PKA (2,500 units/ml) was applied along with ATP and cAMP, robust single-channel currents with ∼4 pA in amplitude were observed (**Fig. 4a**). The PKA-activated currents were outwardly rectifying (**Fig. 4b, c**). In addition, the open probability (Po) of the PKA-activated currents was greater at +60 mV (Po = 0.89) than at -60 mV (Po = 0.14), suggesting a voltage dependence of open probability.

**Fig. 4.**
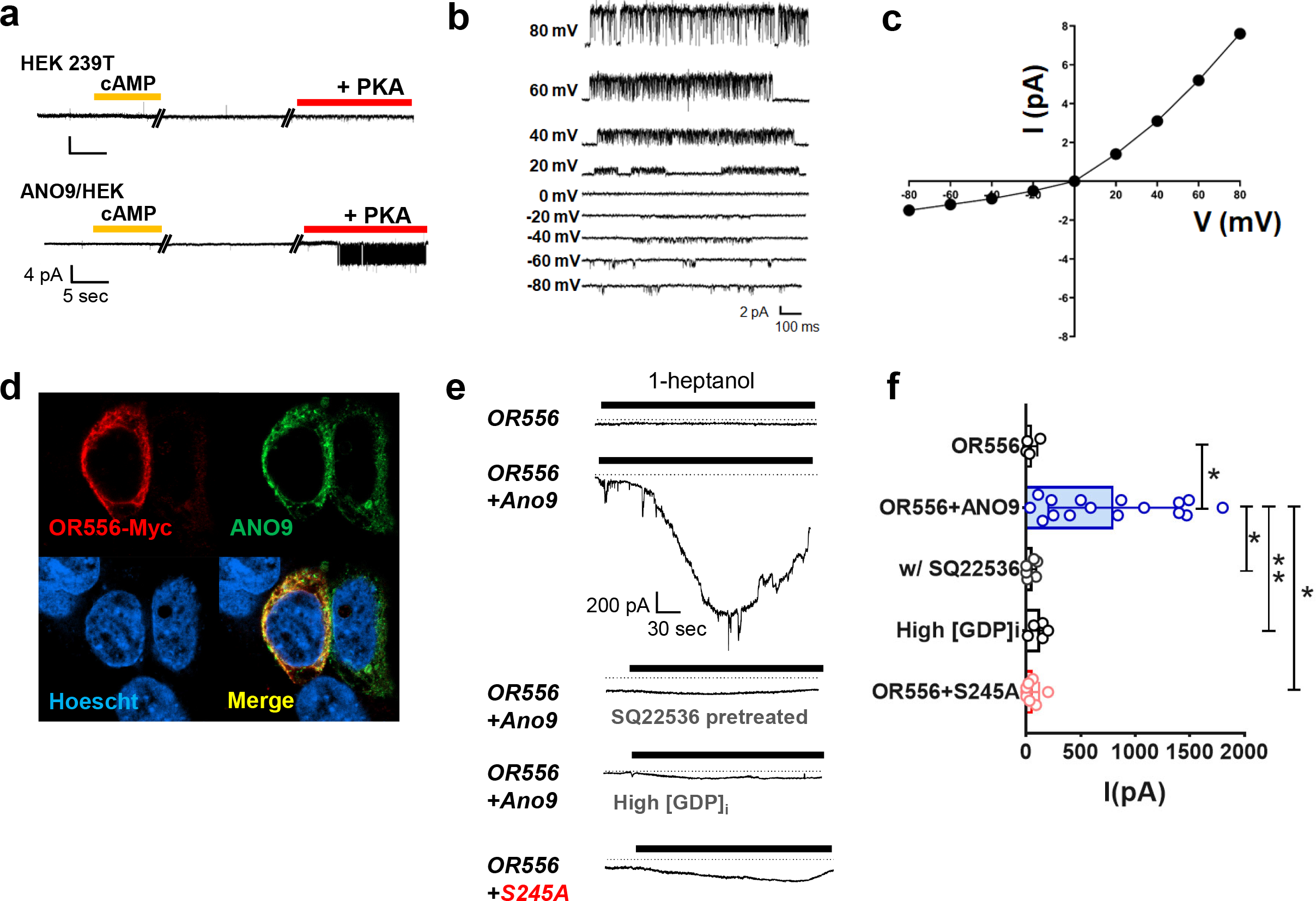
mANO9 is activated by OR stimulation via the cAMP/PKA pathway. **a,** Representative traces of single-channel currents recorded from HEK 293T and ANO9/HEK cells. After forming an inside-out membrane patch, 100 μM cAMP alone or the mixture of cAMP, 2 mM ATP, and purified recombinant PKA (2,500 units/ml) was applied to the bath. Note that ANO9 openings are observed only when the mixture of 100 μM cAMP, ATP, and purified recombinant PKA was applied. Ehold = – 60 mV. **b-c,** The current-voltage (I-V) relationship of single-channel currents of ANO9. Representative traces **(b)** and the I-V curve **(c)** of single-channel currents of ANO9 activated by PKA at various holding potentials. Purified PKA together with ATP and cAMP were applied to an isolated inside-out membrane patch of HEK 293T cells transfected with *Ano9*. **d,** Expression of OR556 and ANO9 in HEK 293T cells identified with Myc antibodies or GFP fluorescence. *Olfr556* (OR556 gene) and *Ano9* were tagged with Myc and eGFP, respectively. **e,** Whole-cell currents of HEK 293T cells transfected with *Olfr556* alone or together with *Ano9*. The bath solution containing 0.1% (v/v) 1-heptanol was applied to the HEK cells. HEK 293T cells transfected with *Olfr556* and *Ano9* pretreated with SQ22536, an inhibitor of adenylate cyclase or 5 mM GDP in the pipette failed to respond to 1- heptanol (3^rd^ and 4^th^ traces, respectively). A HEK 293T cell transfected with *Olfr556* and the *Ano9*/S245A mutant failed to respond to 1-heptanol (5^th^ trace). **f,** Summary of amplitudes of whole-cell currents of HEK 293T cells transfected with *Olfr556* and *Ano9 or* the S245A mutant in different pharmacological conditions. * p < 0.05, ** p < 0.01, One-way ANOVA, Tukey’s post-hoc test.

To identify the PKA phosphorylation site, we performed a mutagenetic approach coupled with electrophysiological experiments. Four putative PKA phosphorylation consensus sites (R-R/K-X-S/T, R-X-X-S/T, and R-X-S/T) were predicted ^34^. These putative sites are Ser85, Ser120, Ser245, and Ser321 residues in mouse ANO9. Each Ser residue was replaced with alanine. Whole-cell currents with different shapes activated by 100 μM cAMP were observed in HEK 293T cells transfected with S85A, S120A, and S321A mutants (**Extended Data Fig. 3a, b**). However, cAMP failed to evoke cation currents in HEK 293T cells transfected with the *Ano9* S245A mutant. Therefore, the Ser245 residue appears to be essential for the activation of ANO9 by PKA.

To determine whether the Ser245 residue is phosphorylated by PKA, we performed an in-vitro phosphorylation experiment coupled with immunoprecipitation ^35, 36^. We overexpressed ANO9 and ANO9-S245A mutant in HEK 293T cells. Cell lysates were precipitated with anti-ANO9 antibodies. We then phosphorylated the immunoprecipitants after incubating them with a purified recombinant catalytic subunit of PKA (2,500 Unit/ml) together with 200 μM ATP. These immunoprecipitants were blotted with anti-phosphoserine antibodies. Western blot analysis showed that the serine phosphorylation level of control ANO9 was higher than that of the ANO9- S245A mutant. (**Extended Data Fig. 5a**). Thus, the phosphorylation at the Ser245 residue of mouse ANO9 is essential for the PKA activation.

**Fig. 5.**
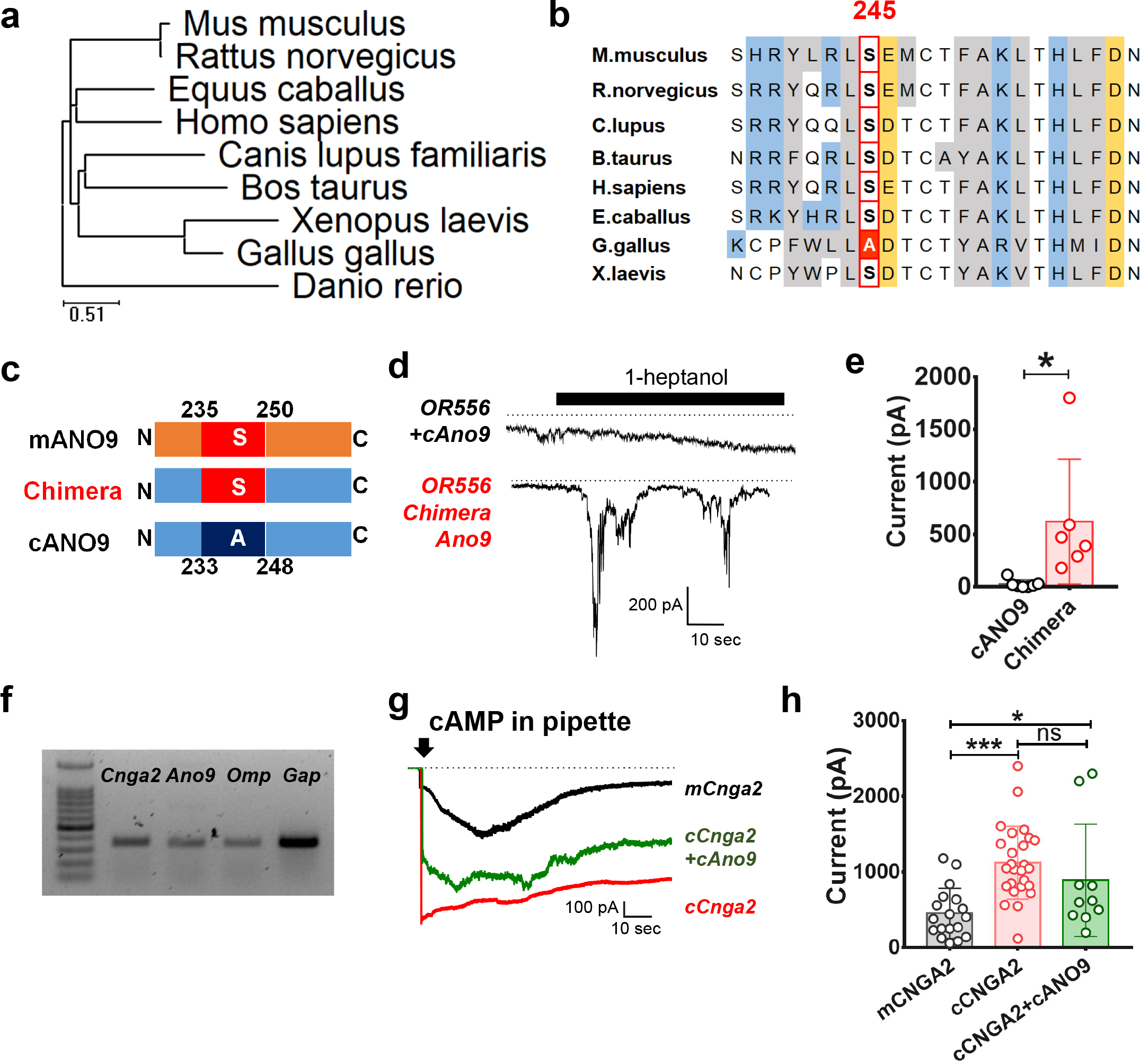
Chick ANO9 is not functional. **a,** Dendrogram based on amino acid sequences among ANO9 orthologs in vertebrates. **b,** Alignment of amino acid sequences of ANO9 orthologs near the Ser245 residue of mouse ANO9 (mANO9). **c,** Cartoon depicting the sequences of mouse, chick ANO9 (cANO9) and a chimera. The chimera was constructed with the cANO9 backbone with an insertion of 16 amino acids of mANO9 flanking the Ser245 residue. **d-e,** Whole-cell current traces **(d)** in response to OR stimulations of HEK 293T cells transfected with c*Ano9* (upper trace) or chimeric *Ano9* (lower trace). Olfr556 was also transfected along with c*Ano9* or chimeric *Ano9*. Summary **(e)** of the amplitudes of the whole-cell currents of each group. * p < 0.05, Student T-test. **f,** RT-PCR products of *Ano9, Cnga2, Omp*, and *Gapdh* (Gap) in the chick OE. **g,** The cAMP-induced currents of mouse *Cnga2*, chick *Cnga2*, and chick *Cnga2*+*Ano9* when expressed in HEK 293T cells. **h,** Summary amplitude of cAMP-induced currents of mouse and chick CNGA2 /HEK and chick CNGA2+mANO9/HEK cells. ns: not significant, * p < 0.05, *** p < 0.001, One-way ANOVA, Tukey’s post-hoc test.

### ANO9 is activated by odorant receptors

We then determined if olfactory receptors (ORs) can stimulate ANO9 in the heterologous expression system. *Ano9* and OR genes were transfected to HEK 293T cells. We selected many ORs (OR43, OR49, OR167, OR168, and OR556) from the OlfactionDB, which is a manually curated database providing comprehensive information about nearly 400 ORs and their respective odorants ^37^. However, among the ORs, only OR556 was expressed in HEK 293T cells (**Fig. 4d, Extended Data Fig. 4a**); ORs are difficult to express heterologously ^38^. Upon whole-cell formation in the HEK 293T cells transfected with *Olfr556* (the OR556 gene) and *Ano9*, robust currents were observed at Ehold of −60 mV when its activating odorant, 1-heptanol, was applied (**Fig. 4e, f**). These 1-heptanol -evoked currents were not observed when *Olfr556* alone was transfected (**Fig. 4e, f**). These 1-heptanol-evoked currents were blocked by the pretreatment of SQ22536, an adenylate cyclase inhibitor, and high GDP (5 mM) in the pipette solution. Furthermore, 1-heptanol failed to evoke a current in HEK293T cells transfected with *Olfr556* and *Ano9*/S245A mutant (**Fig. 4e, f**). These results suggest that ANO9 is activated by OR stimulation via the cAMP/PKA pathway.

### Chick ANO9 is nonfunctional

While it had long been argued whether birds smell, recent studies have demonstrated the bird ability to discriminate odors for feeding, nesting, and other social interactions ^39–41^. We sought to determine if chick olfactory signal transduction is different from mammals. Based on the dendrogram analysis of ANO9 across different species, we found that vertebrate ANO9 orthologs share a gross similarity in amino-acid sequence (51 ∼ 88% sequence identities to mouse ANO9) (**Fig. 5a**). Interestingly, unlike other vertebrates, the residue corresponding to Ser245 of mouse ANO9 essential for the cAMP/PKA activation is substituted with alanine in chick ANO9 (**Fig. 5b**), suggesting that chick ANO9 may not be functional in olfaction. Chick ANO9 has 747 amino acids and shows a 51% sequence identity with mouse ANO9. Upon whole-cell formation in HEK 293T cells transfected with chick *Ano9* and mouse *Olfr556*, 1-heptanol failed to induce currents (**Fig. 5d, e**). We then constructed a chimera that has a chick *Ano9* backbone with an insertion of the 16 amino acids of mouse *Ano9* flanking the Ser245 residue (**Fig. 5c**). The chimeric *Ano9* was activated nicely by the odorant when expressed in HEK293T cells with mouse *Olfr556* (**Fig. 5d, e**). We also checked the functionality of chick CNG channels. Chick and mouse CNGA2 orthologs have an 83% sequence identity. The presence of chick CNGA2 was confirmed with RT-PCR analysis in the chick OE (**Fig. 5f**). In contrast to chick ANO9, chick CNGA2 was activated by intracellular cAMP (**Fig. 5g**). Importantly, the current response of chick CNGA2 to 100 μM cAMP was much greater than that of mouse CNGA2 (**Fig. 5g, h**). The activation kinetics of chick CNGA2 was different from that of mouse CNGA2 as chick CNGA2 was activated rapidly by cAMP (**Fig. 5g**). Also, chick ANO9 failed to amplify the cAMP-induced currents of chick CNGA2 in (**Fig. 5g, h**). Moreover, the chick-mouse chimeric *Ano9* showed a higher Ser-phosphorylation level than that of chick ANO9 (**Extended Data Fig. 5b**). These results suggest that the native chick CNGA2 is functional in olfaction, whereas chick ANO9 is non-functional. Thus, birds appear to have a different mechanism for odorant signal transduction.

### *Ano9* is downregulated in olfactory sensory neurons of Covid-19 patients

Because one of the primary symptoms of Covid-19 is hyposmia or anosmia ^14–17^, we wondered if ANO9 contributes to the anosmia. ANO9 was expressed in the OE in the human nasal cavity (**Fig. 6a**). The ANO9 immunoreactivity was present mainly in the superficial layers of the OE, which overlapped with olfactory marker protein immunofluorescence. We then checked single-cell gene profiling data obtained from healthy control and Covid-19 participants (Chan Zuckerburg initiative single-cell Covid- 19 consortia, http://covid19cellatlas.org) ^42^. Among tissues sampled by the consortium, we analyzed a dataset collected with nasopharyngeal swabs from 37 patients diagnosed with Covid-19 and 21 controls obtained at the University of Mississippi Medical Center. Single-cell RNA sequencing (scRNA-seq) was conducted on 32,588 cells collected from the nasopharyngeal swabs, which annotated 18 cell types (**Fig. 6b**) ^42^. Among the annotated cell types, “Ciliated Cells” were selected for the gene analysis because OSNs have a ciliary region at the apical end. This cell-type cluster contains 4,703 cells from the control and 5,356 cells from the Covid-19 patients (**Fig. 6b**).

**Fig. 6:**
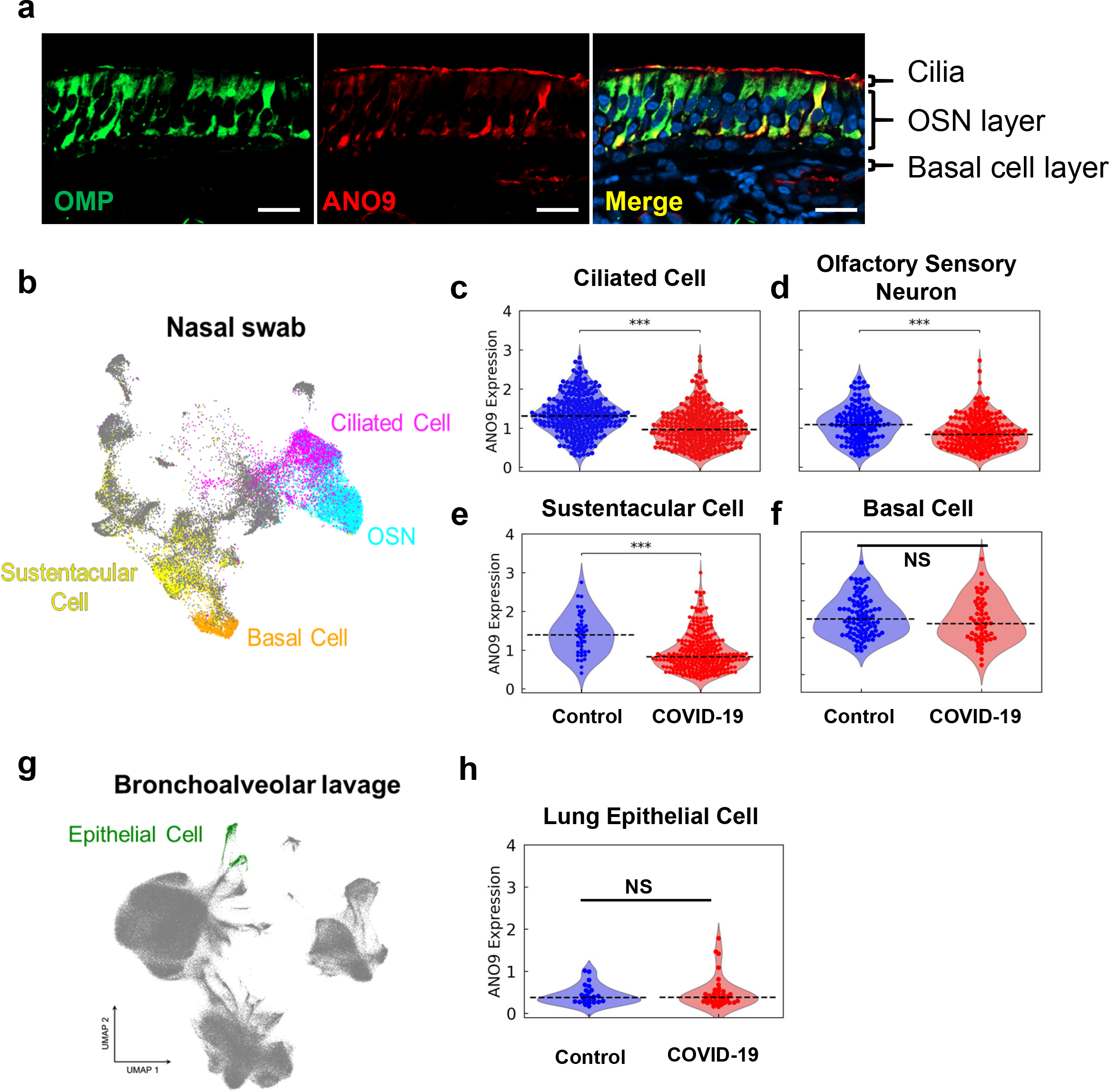
ANO9 is downregulated in nasopharyngeal cells of Covid-19 patients. **a,** Immunofluorescence images of ANO9 in the human OE. Sections of the human OE were stained with anti-ANO9 and OMP antibodies. Nuclei were stained with Hoechst 33342. Scale bars: 20 μm. **b,** Uniform manifold approximation and projection (UMAP) of 32,588 cells obtained with nasal swabs from 37 Covid-19 patients and 21 controls. Eighteen cell types are mapped where ciliated cells (10,059) are highlighted in magenta. Among the ciliated cells, approximately 55.4% can be subcategorized as OSNs (5,573; colored cyan). **c-f,** Distribution of *Ano9* expression (normalized as log(1+UMI per 10K)) in the ciliated cells **(c)**, OSNs **(d)**, sustentacular cells **(e)**, and basal cells **(f)**. Expression level of each gene was obtained from the scRNAseq dataset available online; the Covid-19 Cell Atlas (https://www.covid19cellatlas.org). The boxes, center lines, and whiskers indicate the interquartile range of the non-zero entities, the median values, and the maximum and minimum values within 1.5 × IQR from the box edges, respectively. *** p < 0.001, MAST hurdle model test. **g,** UMAP of 644 cells obtained from 8 Covid-19 patients and 11 controls. Forteen cell types are mapped where epithelial cells are highlighted in green. **h,** Distribution of *Ano9* expression (normalized as log(1+UMI per 10K)) in the lung epithelial cells

However, there was a high number of cells with zero gene expressions due to the limit of scRNA-seq technique, which typically contains between 50 and 99% zeros in the expression matrix ^43^. Thus, we filtered out cells that showed high contents (>15%) of mitochondrially derived genes, assuming these cells were in a poor condition for scRNA-seq analysis ^44, 45^. This left a data set of 3,306 cells × 32,871 genes for COVID-19 patients and 3,000 cells × 32,871 genes for controls. To identify differentially expressed genes relevant to the olfactory signaling, transcript levels of OR genes, ion channels, and other signaling proteins were compared for their differential expression between controls and Covid-19 patients. Among these genes, *Ano9* showed a marked reduction in expression (28.0% reduction, p < 0.001, the MAST hurdle model test) in the Covid-19 patient group compared to that of controls (**Fig. 6c**). However, the annotated ‘Ciliated cells’ also included ciliary cells of the respiratory epithelia. Thus, we narrowed down the putative OSNs. We counted cells positive for the *Gnal* or *Stoml3* gene, representative marker genes for mature OSNs ^46^. These cell clusters contained 1,661 and 2,196 cells for control and Covid-19 patient groups, respectively. In this cell cluster, *Ano9* was significantly reduced in the Covid-19 patient group (23.0%, p < 0.001, the MAST hurdle model test) (**Fig. 6d**). Other genes such as *Cnga4*, *Cngb1*, and *Prkaca* were significantly increased, whereas *Gnal* was significantly decreased in the patient’s group (**Table I**). The number of cells with *Ano2*, *Cnga2, Prkacg,* and individual OR genes were too small to count for statistical comparison. In this data set, we also compared the levels of Ano9 in other cell types such as “Sustentacular cells” and “Basal Cells”. Sustentacular cells are known to support and protect OSNs in the OE, whereas basal cells are precursor cells for sustentacular or OSNs ^47^. Sustentacular cells were defined if cells excluded from basal cells, blood-borne cells, and OSNs were positive with Sox2 and CYP2A13 genes, marker genes for sustentacular cells ^46^. Sustentacular cell clusters contained 282 and 2083 cells for control and Covid-19 patient groups, respectively. As shown in **Fig. 6e**, the *Ano9* mRNA level in the sustentacular-cell cluster from the Covid-19 patient group was significantly lower than that of the control group, whereas the *Ano9* mRNA level in the basal-cell cluster was comparable in both groups (**Fig. 6e, f**).

**Table I:**
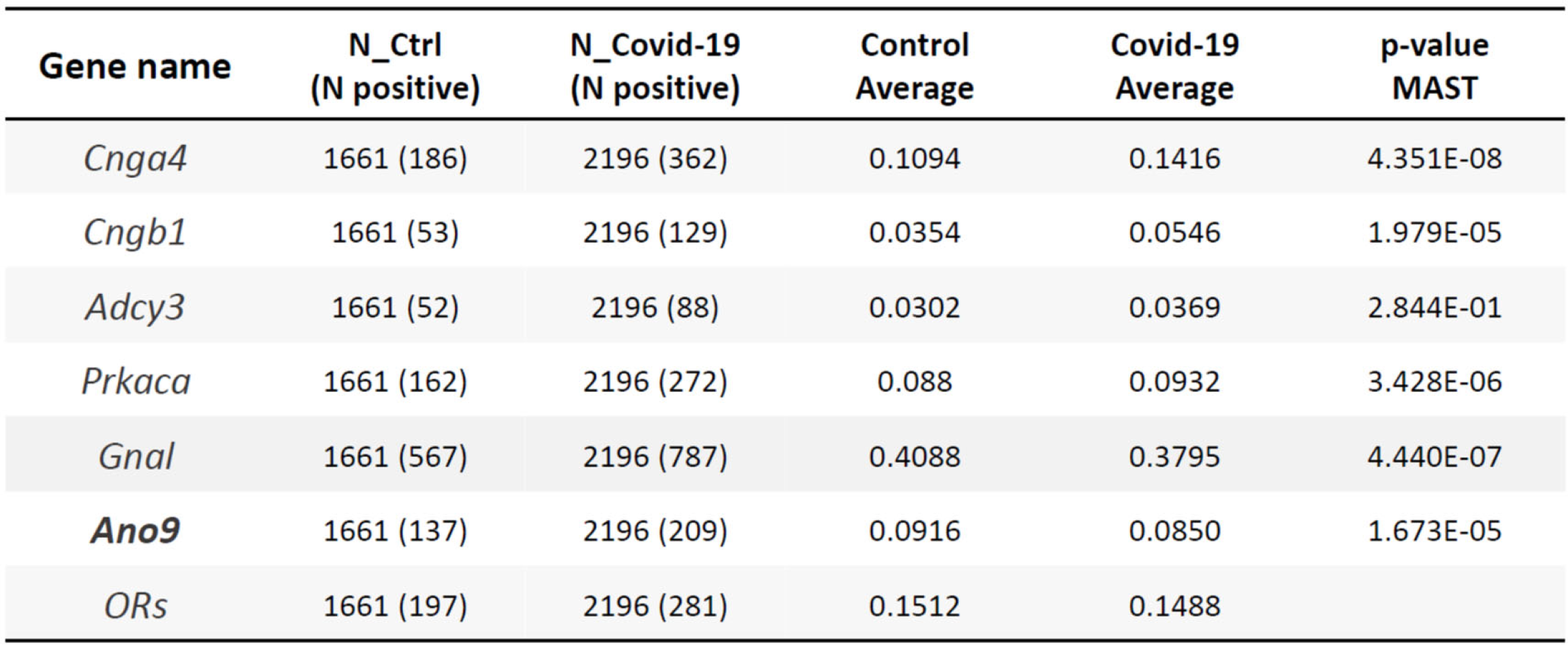
Statistical tests of differential expressions of olfactory signal genes in the putative OSN cluster of Covid-19 patients and healthy controls. **Differential** expression of ion channels and signaling proteins in the OSN cluster between the SARS-CoV-2 positive and control groups are summarized. OSNs were selected if the ‘Ciliated cells’ had a positive value of *Gna*l or *Stoml3*. Zero values were included for all genes. Numbers of positive cells, the average transcription level of each gene (for normalization; log(1+UMI per 10K)) and *p*-values from the MAST hurdle model test are shown (see **Methods**).

| **Gene name** | **N_Ctrl**<br>**(N positive)** | **N_Covid-19**<br>**(N positive)** | **Control**<br>**Average** | **Covid-19**<br>**Average** | **p-value**<br>**MAST** |
| --- | --- | --- | --- | --- | --- |
| *Cnga4* | 1661 (186) | 2196 (362) | 0.1094 | 0.1416 | 4.351E-08 |
| *Cngb1* | 1661 (53) | 2196 (129) | 0.0354 | 0.0546 | 1.979E-05 |
| *Adcy3* | 1661 (52) | 2196 (88) | 0.0302 | 0.0369 | 2.844E-01 |
| *Prkaca* | 1661 (162) | 2196 (272) | 0.088 | 0.0932 | 3.428E-06 |
| *Gnal* | 1661 (567) | 2196 (787) | 0.4088 | 0.3795 | 4.440E-07 |
| **Ano9** | 1661 (137) | 2196 (209) | 0.0916 | 0.0850 | 1.673E-05 |
| *ORs* | 1661 (197) | 2196 (281) | 0.1512 | 0.1488 |  |

To test whether this reduction of *Ano9* transcripts in Covid-19 patients is unique to ciliated cells in the nasopharyngeal epithelium, we profiled another scRNAseq data from lung epithelial cells in bronchoalveolar lavage fluid (BALF) in the Covid-19 cell atlas consortium. The study population consisted of COVID-19 patients (n = 8) and controls (n = 11) recruited from the Ghent University Hospital. The control group was a combination of non-SARS-CoV-2 respiratory disease (n = 9) and healthy subjects (n = 2). Among the 14 annotated cell types, we chose the “Epithelial cell” cluster (**Fig. 6g**).

In this cell cluster, we filtered out cells that failed to show positive signals for epithelial cell housekeeping genes, GAPDH, TUBB4B, or TUBB ^48^, which led to 388 and 256 cells for patient and control groups, respectively. As shown in **Fig. 6h**, the *Ano9* transcript level in Covid-19 patient group did not differ from that of control group

## DISCUSSION

Odorants stimulate ORs that elevate cAMP levels in OSNs via adenylate cyclase activity ^49–51^. The cAMP, in turn, opens CNG channels. Olfactory CNG channels in OSNs consist of three subunits: CNGA2, CNGA4, and CNGB1 ^32, 52, 53^ with a stoichiometry of two CNGA2, a CNGA4, and a CNGB1b ^30^. Genetic disruption of *Cnga2* induces almost complete loss of EOG responses to odorants ^6^, implying that CNG channels are considered the main transduction channel in olfaction ^6, 54^. However, CNG channels desensitize rapidly as Ca^2+^ binds to CNG channels limiting the Ca^2+^ influx in the OE ^55^. Because ANO9 is present in OSNs and activated by intracellular cAMP, odorant-induced cAMP triggers ANO9 in OSNs. Furthermore, intracellular Ca^2+^ and the depolarization induced by the CNG channel opening augments the ANO9 activity. Thus, positive feedback cycles for opening ANO9 will occur once triggered by the CNG channel activation, amplifying the sensory signals in OSNs (**Extended Data Fig. 6**). The present study suggests that ANO9 is a candidate for the alternative cAMP-induced Ca^2+^ signals in OSNs. Firstly, i) ANO9 was expressed in the OE, ii) odorants activated ANO9 in a heterologous system, and iii) it amplified the cAMP-evoked CNG channel currents. Secondly, the genetic ablation of *Ano9* reduced cAMP-induced currents in isolated OSNs, EOG signals to odorants, *in-vivo* neural activity in the OB, and sensitivity to odorant discrimination. Also, we found that chick ANO9 is nonfunctional, whereas chick CNGA2 shows a greater response to cAMP. Therefore, it is likely that mammals require signal amplification by ANO9 for olfaction, whereas birds do not. Thus, the present study provides molecular insights into the diversity in olfactory transduction signaling between birds and other vertebrates.

### Anoctamins and olfactory transduction

Anoctamins (TMEM16A-K) are a gene family with numerous physiological functions. ANO1 was cloned first in the family as a Ca^2+^-activated Cl^-^ channel ^29, 56, 57^. Anoctamins consist of 10 homologs from ANO1/TMEM16A to ANO10/TMEM16K, which show multiple modes of actions. ANO1 and ANO2 are Cl^-^ channels activated by Ca^2+^ that mediate numerous physiological functions from transepithelial fluid secretion to nociception ^29, 58, 59^. ANO5 and ANO6 are scramblases that neutralize polarized phospholipids ^60–62^. Adding to the diverse actions of Anoctamins, ANO9 is a cation channel activated by phosphorylation by PKA. Despite various modes of action, the Ca^2+^ dependency appears common to all Anoctamins as ANO1 and ANO2 are activated by Ca^2+^, and the ANO6 scramblase activity also requires Ca^2+ 62, 63^. ANO4 is also a calcium-dependent cation channel (Reichhart et al., 2019). Thus, most of Anoctamin family proteins appear to require intracellular Ca2+ for their activation or as co-factor for augmenting their activity. It is not surprising that ANO9 is augmented with intracellular Ca^2+^ (**Fig. 3g, h**) ^33^.

The role of Ca^2+^-activated Cl^-^ channels in olfaction had been studied extensively ^64–66^. Because the level of intracellular Cl^-^ is high in OSNs ^67^, Cl^-^ channels’ opening leads to depolarization in OSNs ^64^. Indeed, ANO2 is expressed in OE ^10, 64, 68^. A comparison of the biophysical properties of native olfactory Cl^-^ currents in OSNs and ANO2 revealed a remarkable similarity ^69^. Despite the role of native Ca^2+^-activated Cl^-^ channels in OE, *Ano2*-deficient mice are normal in olfactory behavior tasks ^10^.

### Olfaction in birds

It had been considered for a long time that birds do not smell. However, numerous reports now show that birds smell odorants for feeding, reproduction, and social interactions ^39–41^. The gross anatomy of the bird’s olfactory system does not differ much from those of amphibians, reptiles, or mammals ^70^. In addition, chick OSNs possess a similar morphology to those of other vertebrates ^71^. Birds have a large number of OR genes, suggesting an active role of olfaction in birds ^72, 73^. However, the molecular mechanisms underlying the chick olfactory signal transduction are largely unknown. Surprisingly, chick ANO9 was non-functional, whereas chick CNGA2 was more active than mouse CNGA2. These results imply that olfactory transduction in birds is largely dependent on CNG channels only, which is a stark contrast with the mammalian olfactory system.

### Covid-19 related anosmia

One of the symptoms of Covid-19 patients is a loss of chemical senses, including smell. However, there are no etiological explanations for this except for possible inflammation caused by SARS-CoV2 infection ^74^. A noticeable reduction in the *Ano9* transcript level in OSNs and sustentacular cells, but not in the lung epithelial cells, was observed in Covid-19 patients (**Fig. 6d, e, h**). Numerous studies report a high incidence of anosmia among Covid-19 patients. Depending on the ethnicity and condition of the studies, the prevalence of olfactory dysfunction in Covid-19 patients varies from 47.9 to 85.6% ^14, 17, 75^. Thus, about two-thirds of Covid-19 patients suffer from anosmia. Even though about a third of Covid-19 patients may not be anosmic, we still see a reduction in *Ano9* mRNA level in OSNs of the Covid-19 patients. The change in the *Ano9* expression may be greater than observed in this study If the *Ano9* mRNA level is checked only in the anosmic patients. This downregulation of *Ano9* may contribute to the Covid-19-associated anosmia. However, the route and underlying mechanisms for the downregulation are not known. A subset of supporting cells in the human OE, but not OSNs, express ACE2, the SARS-CoV2 Spike protein receptor ^47^, arguing against direct infection of the coronavirus in OSNs. However, low-level expression of ACE2 can induce SARS-CoV2 entry ^76^. Or, paracrine effects from virus-infected supporting cells would account for the downregulation of *Ano9* ^45, 47^.

In summary, ANO9 expressed in the OE augments the odorant-induced olfactory signals via the cAMP/PKA pathway. The amplifying action of ANO9 contributes to olfactory function because ANO9-deficient mice showed impairment of olfaction. In addition, ANO9 may be a risk factor of Covid-19-associated anosmia because Ano9 mRNA level is substantially reduced in nasopharyngeal epithelial cells of Covid-19 patients. These results now suggest that ANO9 actively participates in the olfactory signaling in the OE.

## MATERIALS AND METHODS

### Human Samples

Two tissue samples of the human OE were acquired from two male cadavers (age: 68 years and 80 years). Due to the domestic law that cadavers must be donated and managed under informed consent and ethical process, the institutional review board (IRB) of The Catholic University of Korea has decreed that studies involving cadavers do not need to be reviewed. All cadavers stored at Catholic Institute for Applied Anatomy qualified as materials for education and research according to domestic law. The Institutional Steering Committee approved cadaver use for this research on March 10, 2021 (Approval number: R21-A005).

### Animals

Animal use was approved by the Institute of Laboratory Animal Resources of the Korea Institute of Science and Technology. Adult male and female C57BL/6 background, transgenic, and *Ano9* KO mice were used for electrophysiological recordings, *in-vivo* Ca^2+^ imaging, or behavioral tests. In standard housing conditions, mice were housed in groups (two to five mice per cage) in an open-top cage system. Water and food were available *ad libitum*. All animals were kept at light and dark in a 12-hour cycle.

For the generation of *Ano9^fl/fl^* mice, two *loxP* sites were introduced within the ANO9 locus by using the CRISPR/Cas9 system. The first and second *loxP*s were inserted to introns 7-8 and 9-10, respectively. Cas9 nuclease, validated sgRNAs, and donor template containing *loxP* sites were co-injected into single-cell C57BL/6 zygotes. The *loxP* regions of the mouse were confirmed after genotyping DNAs extracted from the mouse tail. To delete *Ano9* in the whole body, the mice genotyped for the presence of the *loxP* sites (*Ano9^fl/fl^*) were crossed with B6.C-Tg(CMV-cre)1Cgn/J (CMV-cre) to produce CMV-cre;*Ano9*^fl/-^ mice. The CMV-cre;*Ano9*^fl/-^ mice were crossed with *Ano9*^fl/fl^ mice to generate *Ano9* deficient mice (CMV-cre;A*no9*^fl/fl^).

### Odor preference test

This test was designed to identify the specific ability to sense attractive or aversive scents ^21^. To habituate, each mouse was placed in a cage for 30 min. After the habituation, the mouse was transferred to a test cage where a dish containing water with a multi-perforated lid was placed. Exploring time to the dish during the 3 min test period was measured. After changing the dish containing 10% (w/v) peanut butter or 2- methylbutyric acid (193070, Sigma) solution, the exploration time to the dish was also measured for 3 minutes. The mouse behavior was recorded with a video camera for later analysis.

### Odor discrimination test

The odor discrimination test was performed as described previously ^22, 23^. Briefly, mice were trained for six days to associate (+)-carvone (818410, Sigma) with a sugar reward in a restricted feeding state. Mice were placed in a cage with (+)-carvone paired with the sugar reward and transferred to a cage with a (-)-carvone (124931, Sigma) unpaired with the sugar reward for 20 min daily. During the training, (+)-carvone (20 μl) and (-)-carvone (20 μl) were put in a polystyrene dish (35 x 10 mm) with a multiperforated lid and placed on the bedding in the center of the cage (16 cm height, 22 cm width, 30.5 cm length). Several pieces of crystal sugar were placed on the lid of the dish containing (+)-carvone as a reward. On day 5, the (+)-carvone and (-)-carvone odorants were placed separately under the bedding (5 cm depth) at either end of the cage. At this time, sugar was not present on the (+)-carvone-containing dish. Mice behavior was recorded on a video camera to measure the time spent digging for each side during the 5-minute test. On days 6 and 7, only one odorant was placed under the bedding on one side of the cage without sugar (day 6 (+)-carvone; day 7 (-)-carvone). Mouse digging behavior was also recorded for each test. On days 5 and 6, training was also performed after the test.

### *In-vivo* Ca^2+^ imaging of the olfactory bulb

Surgery and *in-vivo* imaging were described in a previous report ^53^. Briefly, male and female adult mice (9-12 weeks old) were anesthetized with intraperitoneal injection of Ketamine/Xylazine (90 mg/10 mg/kg). Anesthesia was maintained by additional injections of Ketamine/Xylazine (45 mg/5 mg/kg). The animal body temperature was maintained at ∼36°C using a heating pad. Craniotomy was carried out to expose both olfactory bulbs. Bupivacaine (0.5%) was injected prior to a longitudinal incision from behind the ear to the anterior part of the olfactory bulb. The skin was retracted to expose the skull. A craniotomy was carried out to expose both olfactory bulbs. A head post was attached to the skull with cyanoacrylic glue and dental cement. The skull above the two hemibulbs was thinned with a dental drill. After the craniotomy, the olfactory bulb was covered with Ringer’s solution.

Oregon Green 488 BAPTA-1, AM (Thermofisher Scientific) was freshly dissolved in 20% Pluronic F-127 in DMSO and diluted in a solution (in mM: 150 NaCl, 2.5 KCl, and 10 HEPES, pH 7.4) containing 2.5% Fast Green (Sigma) to produce 0.5 mM Oregon Green. The final Pluronic F-127 concentration was ≈ 0.5%. The dye solution was sonicated and filtered (0.45 µm Ultrafree filter, Millipore). The dye (1 ml) was injected into the glomeruli layer of the olfactory bulbs through a glass microelectrode with a tip diameter of 10 -12 um using a Nanoinject III (Drummond). Fluorescence imaging was performed one hour after the injection of the dye solution. The olfactory bulbs were covered with 2% agarose in Ringer’s solution, and a glass coverslip was gently pressed onto the agarose to suppress the brain movement. Odorants were delivered with an olfactometer four times with 1 min-intervals. Odorants (10%) were diluted from saturated vapor with humidified air. Either 1-heptanol or a mixture of odorants (2-heptanone, (+)- carvone, and (-)-citronellal (Sigma-Aldrich)) was used.

Optical imaging of the olfactory bulbs was performed using the wide-field section of a modified Sutter MOM (Movable Objective Microscope, Sutter Instruments) equipped with a NeuroCCD SM-256 camera (RedShirtImaging). The light from a Prizmatix LED (UHP-T-LED-460-LN) was used with a Nikon, 10x. 0.45 NA lens using a 472/30 nm band-pass excitation filter (Semrock FF02-472/30), a 495 nm dichroic mirror, and a 496 nm long-pass emission filter (Semrock FF01-496/LP). The emitted fluorescence was recorded at 125 frames/second using NeuroPlex software (RedShirtImaging, Decatur, GA). Imaging data were analyzed with NeuroPlex. The location of calcium signals in the image was determined with the Frame Subtraction function. The temporal average of the frames 1-2 s preceding the odorant stimulus was subtracted from a 1-2 s temporal average around the response peak. The resulting activation maps indicated the pixels with fluorescence intensity changes (**Fig. 2c, d**).

The calcium signal traces are shown as fractional fluorescence changes by dividing the fluorescence intensity change in the glomerulus by the fluorescence intensity preceding the stimulus.

### EOG recordings

Organ preparation for the EOG recordings was described elsewhere ^3, 6^. Briefly, the exsanguination was performed through an open heart immediately after deep anesthetization to minimize blood in the nasal tissue. The mouse head was excised longitudinally along the midline to expose the olfactory turbinates. The half head was mounted in a recording chamber with 2% agarose gel. Odorants were freshly diluted to make 0.1 or 10 mM in Ringer’s saline (in mM, 145 NaCl, 5 KCl, 10 HEPES, 2 MgCl2, and 2 CaCl2, pH 7.2) and mixed with vigorous vortexing. Odorants were delivered to the turbinate through a computer-controlled air injection system (HSPC-2-SB, ALA Scientific Instruments) with an approximate flow rate of 0.2 ml/sec. The reference Ag/AgCl electrode was placed in the bath. A glass pipette electrode for recording was pulled with a micropipette puller (P-97, Sutter Instruments) followed by polishing the tip of the pipette with a microforge (MF-830, Narishige, Japan). The glass pipette was filled with the 0.9%-agar containing Ringer’s saline. Evoked extracellular voltage was recorded from the surface of turbinates IIb or III and amplified with a differential amplifier (DP-304, Warner Instruments). The output signals of the amplifier were filtered at 10 kHz. The electrical signals were digitized at 500 Hz and analyzed using pClamp 10.

### Gene cloning and mutation

All mutants were generated from a mouse *Ano9* construct. The amino acid substitution methods were used to construct *Ano9* mutants using a site-directed mutagenesis kit (Muta-Direct™, Intron Biotechnology). The construction of mutants was verified with DNA sequencing. The chick *Ano9* construct (XM_015286257.2) was chemically synthesized. The chick and mouse *Ano9* chimera mutant was cloned using an overlap extension PCR strategy. We subcloned chick *Ano9* into pEGFP-N1 vector from the XM_015286257.2:1133-3649 PREDICTED: *Gallus gallus* anoctamin 9 (*Ano9*), transcript variant X3, mRNA. The chimeric sequence was replaced between 696 and 743 of *Gallus gallus Ano9* by the *Mus musculus Ano9* sequence, -GGC GAC CAC AGT CAC AGA TAC CTG CGA CTC TCA GAG ATG TGC ACT TTC- (NM_178381.3). All mutant or original channel genes were tagged with eGFP or subcloned to a transfection vector, pEGFP-N1.

### Cell culture

HEK 293T cells were used for transfections of mouse or chick *Ano9, Cnga2,* and various GPCRs. The transfected HEK 293T cells were incubated at 5% CO2 at 37^◦^C in the Dulbecco’s modified Eagle’s medium supplemented with 10% fetal bovine serum, 10 units/ml penicillin, and 10 μg/ml streptomycin. The HEK 293T cells were mixed with 1 μg of mouse *Ano9* cDNA (m*Ano9*-pEGFP-N1) and 3 μl FuGene HD (Roche Diagnostics). The transfected cells were plated onto glass coverslips that were kept in a 35-mm round Petri dish. The transfected HEK 293T cells were used for current recording for 24 to 48 h after the transfection.

### Primary cultures of OSNs

Dissociated OSNs were prepared as previously described ^77^. After decapitation, the skin overlying the skull was peeled away. The lower part of the head was cut along the jaw line and removed. The upper front teeth were also removed by a coronal cut parallel to the teeth. The skull was cut along the midline of the nose. From two hemiheads, the nasal septum and olfactory turbinates were pulled off. The OE was isolated from nasal septum and the attached parts of turbinates. The separated OE was minced by scissor, and incubated in Ca^2+^ free Na-HEPES solution supplemented with 2 units/ml papain (P4762, Sigma) and 200 μM L-Cysteine (C7352, Sigma) for 10 min. For stopping the enzyme activity, cells were transferred to Ringer’s solution containing 400 μg/ml leupeptin (L8511, Sigma), 100 μg/ml bovine serum albumin, and 20 μg/ml DNase 1 followed by a gentle trituration with a flame-polished Pasteur pipette. Cells were centrifuged at 300×g (1,500 rpm) for 3 min. After the supernatant was removed, the pelleted cells were diluted with 1 ml of Ringer’s solution. Cells were allowed to settle onto glass coverslips coated with 5 mg/ml concanavalin-A (C2010, Sigma) and 5 mg/ml poly L-lysine for 5 min. The OSNs were stored at 4 °C for up to 5 hours for later experiments.

### Antibody Production

A peptide spanning the N terminus of mouse ANO9 (72∼92 amino acid: KDQKKVFFGIRADSDVIDKYR-C: C-term cysteine was added for conjugation efficiency) was synthesized. KLH/BSA conjugation was performed for effective immunization and generation of anti-peptide antibodies. New Zealand White Rabbits were immunized with 500 μg of the peptides in an emulsion with complete Freund’s adjuvant. After 4 weeks, Rabbits were then boosted three times at 2-week intervals with 200 μg of the peptides in an emulsion with incomplete Freund’s adjuvant. After the 3^rd^ boosting, rabbits were sacrificed, and serum were collected to confirm antigen specificity by ELISA test. The final serum was used for immunofluorescence analysis (AbFrontier, South Korea).

### Immunofluorescence

For the staining of sections of the OE, the tissues were fixed by immersion in 4% paraformaldehyde solution overnight at 4°C, cryoprotected with 30% sucrose in phosphate-buffered saline (PBS) overnight, embedded in Tissue-Tek® O.C.T. Compound (Sakura Finetek), and frozen. The nasal bone was decalcified after the immersion of sections in Calci-Clear^TM^ Rapid (National Diagnostics) for 1 hr after fixation. Cryosections of 20 µm thickness were placed on silane-coating slides (Muto Pure Chemicals). After O.C.T compound was rinsed in PBS buffer, for the permeabilization, the sections were submerged in PBS containing 0.2% Triton X-100 for 10 minutes. The sections were incubated with PBS-buffered blocking solution containing 0.1% Triton X- 100 and 1% bovine serum albumin for 1 hr. Mouse polyclonal anti-ANO9 antibody (1:100, AbFrontier, South Korea), monoclonal anti-c-Fos antibody (1:1000, ab208942, Abcam), Tuj-1 (1:500, T5076, Sigma), Ac-Tub (1:500, T7451, Sigma), or olfactory marker protein (1:200, 019-22291, FUJIFILM Wako Chemicals) was diluted in 0.1% Triton X-100 in 1 x PBS containing 1% bovine serum albumin. The sections were incubated with the diluted primary antibody overnight at 4°C. After three 5-min rinses in PBS, the sections were incubated with the secondary antibody conjugated to Alexa 488 or 594(1:1000, A-21467, A-21442, Thermo Fisher Scientific) for 2 hours at room temperature. After washing three times for 5 min each, the sections were incubated with Hoechst 33342 (1:2000, H3570, Thermo Fisher Scientific) for 10 minutes at room temperature. Labeled sections were imaged with an LSM800 confocal microscope (Carl Zeiss).

For determining the expression of c-Fos in the olfactory bulb, mice of both genotypes were exposed to an odorant mixture including 1-heptanol, 2-heptanone, (-)- citronellal, (+)-limonene, and (-)-carvone (diluted 1∶100 in water, Sigma). A piece of filter paper (2 cm × 4 cm) was soaked with the odorant mixture. Each mouse was placed in a cage where the filter paper was placed in the center. After the 30-min exposure, the mice were sacrificed for c-Fos immunofluorescence preparation. Mice without the filter paper were grouped as the naïve control.

### Channel current recordings

Whole-cell current recordings were obtained using the voltage-clamp technique with an Axopatch 200B amplifier (Molecular Devices) as descried previously ^33, 78^. Briefly, whole-cell currents were measured after breaking the plasma membrane under the pipette tip. The resistance of the glass pipettes was about 3 mΩ. The junctional potentials were cancelled to zero. The bath solution contained (in mM) 140 NaCl, 5 KCl, 2 CaCl2, 2 MgCl2 and 10 NaOH-HEPES adjusted to pH 7.2. The pipette solution contained (in mM) 140 KCl, 2 MgCl2, and 10 KOH-HEPES adjusted to pH 7.2. Ca^2+^ was chelated with 5 mM EGTA to make 1.0 μM free Ca^2+^ in the pipette solution. The free Ca^2+^ was calculated using WEBMAXC (https://somapp.ucdmc.ucdavis.edu/pharmacology/bers/maxchelator/webmaxc/webmaxcS.htm). Ehold = -60 mV.

For single channel current recording, the pipette tips were insulated with Sylgard resin (Dow Corning #184) to reduce electrical noise ^79^. Tip resistance was ∼5 mΩ. After a giga-seal was formed, the glass pipette was pulled quickly from the cell to make an inside-out membrane patch. The pipette solution contained (in mM) 140 CsCl, 2 CaCl2, 2 MgCl2 and 10 CsOH-HEPES adjusted to pH 7.2. The bath solution contained 140 CsCl, 2 MgCl2 and 10 CsOH-HEPES adjusted to pH 7.2. To apply purified PKA proteins, purified recombinant PKA proteins (2,500 units/ml, New England Biolabs) along with 100 μM cAMP, 1 mM ATP were added to the bath solution.

The channel open probability (*P_o_*) and amplitudes of single-channel currents were analyzed with pCLAMP software. *P_o_* of single channels was estimated from the ratio of the areas under the curves representing open events divided by the sum of the areas under the curves representing open and closed events. Holding potential was set at -60 mV unless stated otherwise. The output of the amplifier was fed to an analog/digital converter (Digidata 1440, Molecular Devices) and stored in a personal computer. The pClamp 10 software was used for the I-V curves and other biophysical analysis.

### Dendrogram

Vertebrate ANO9 amino-acid sequences with the following accession numbers were acquired from NCBI reference sequence: NP_001012302.2 (*Homo sapiens*), NP_848468.2 (*Mus musculus*), XP_006230651.1 (*Rattus norvegicus*), XP_022260989.1 (*Canis lupus familiaris*), XP_015141743.1 (*Gallus gallus*), XP_023510617.1 (*Equus caballus*), XP_010819602.2 (*Bos Taurus*), XP_018112961.1 (*Xenopus laevis*), and XP_001921968.4 (*Danio rerio*). Multiple sequence alignment was obtained using the CLUSTAL OMEGA program (https://www.ebi.ac.uk/Tools/msa/clustalo/) to compare the ANO9 sequences of various species. The dendrogram was drawn by the Mega-X program (https://www.megasoftware.net/).

### In-vitro phosphorylation experiment

For the immunoprecipitation assay, HEK 293T cells were transfected with the mouse Ano9, S245A-Ano9, chick Ano9 or chick-mouse chimeric Ano9. The transfected cells were washed with ice-cold phosphate-buffered saline. The cells were harvested with RIPA buffer supplemented with a protease inhibitor cocktail (cOmplete^TM^, Roche) and a phosphatase inhibitor cocktail (Sigma, P0044). The cells were lysed with repeated vortexing for 30 min. The cell lysates were mixed with anti-ANO9 antibodies and incubated overnight in 4°C. The immune complexes were collected after binding to protein A beads and washed twice with 1X kinase buffer. Pellets were suspended in 40 ul 1X kinase buffer supplemented with 200 μM ATP and the recombinant catalytic subunit of PKA. Suspended lysates were incubated 30 miniates at 30°C. The samples were heated at 95°C for 5 minutes with 2X SDS sample buffer. The samples in SDS buffer were separated by SDS-PAGE. Then, these immunoprecipitants were blotted with anti-phosphoserine antibodies

### Statistical analysis

All results are expressed as means ± standard errors (S.E.). The statistical significance of the difference in means was determined by a one-way analysis of variance (ANOVA) followed by Tukey’s post-hoc test or two-way ANOVA followed by Bonferroni’s post-hoc test for multiple comparisons. The comparison of two means was made by the Student T-test. Statistical significance was accepted at p values of less than 0.05.

### scRNA-seq Data analysis

The procedures for recruiting participants, collecting samples, sequencing mRNAs, and analysis of genes were described previously ^42^. Briefly, participants were recruited from outpatient clinics, medical surgical units, intensive care units, or endoscopy units at the University of Mississippi Medical Center (UMMC). The UMMC Institutional Review Board approved the study under IRB#2020-0065. Participants included 35 SARS-CoV- 2 positive individuals (19 male and 16 female), 21 SARS-CoV-2 negative individuals (11 male and 10 female). The scRNA-seq was performed by the Shalek’s lab (SeqWell S3 Master Protocol (http://shaleklab.com/wp-content/uploads/2019/07/SeqWell-S3-Protocol.pdf). The data for lung epithelial cells in bronchoalveolar lavage fluid (BALF) was collected from the Ghent University Hospital. The Ethics Committee of Ghent University Hospital (Belgium), AZ Jan Palfijn (Belgium) and AZ Maria Middelares (Belgium) approved the study under G0G4520N. The cohort consisted of 8 COVID-19 patients and 11 control cases as a combination of 9 non-SARS-CoV-2 respiratory disease and 2 healthy individuals. The scRNA-seq was conducted at the VIB Nucleomics Core (VIB, Leuven, Belgium).

In both data sets, the male to female ratios were about 1:1. The full data sets are available on the Covid-19 Cell Atlas (https://www.covid19cellatlas.org), and the details of recruiting patients/controls, data collections, and scRNA-seq methods are described in the preprint ^42^.

### Statistical analysis of the scRNA-seq data

Shalek and colleagues had clustered and annotated the sequenced cells into 18 cell types ^42, 45^. Among the cell types, “Ciliated Cells” were the most highly populated and showed positive OR gene expression. Therefore, we limited our analysis to this type. Six SARS-CoV-2 negative individuals who had respiratory symptoms were excluded from the control group; this left fifteen healthy control subjects (6 male and 9 female). Thirty-five SARS-CoV-2 positive individuals were compared with the control group. We further performed quality control by excluding cells with greater than 15% mitochondrial DNA reads. This resulted in 3,306 cells × 32,871 genes for patients and 3,000 cells × 32,871 genes for controls.

Three genes in the OE (*Ano9, Cnga4,* and *Cngb1*) and three olfactory signaling proteins (*Gnal, Adcy3, and Prkaca*) were chosen for the differential expression analysis. The analysis between the groups was determined with the MAST hurdle model test ^80^, a two-sided test for the null hypothesis that two independent samples have identical expected values. The test was shown to be reliable and robust for scRNA-seq analysis ^81^. The analysis was performed with tailored Python and R scripts utilizing the following packages: Scanpy (v.1.7.1) ^82^ was used to import data and SciPy (v.1.4.1) for t-test ^83^ (see Code availability). Uniform manifold approximation and projection (UMAP) was performed using the cellxgene program (v0.16.7) (http://shaleklab.com/wp-content/uploads/2019/07/SeqWell-S3-Protocol.pdf).

### Code Availability

Codes are available fro the authors upon reasonable request.

## ACKNOWLEDGMENTS

We are very grateful to Prof. Alex Shalek and the members of his laboratory at MIT for their technical information on single cell RNA sequencing datasets of Covid-19 patients and healthy controls. This study was supported by the National Research Foundation of Korea (2020R1A3A300192911).

---

**Extended Data Fig. 1.**
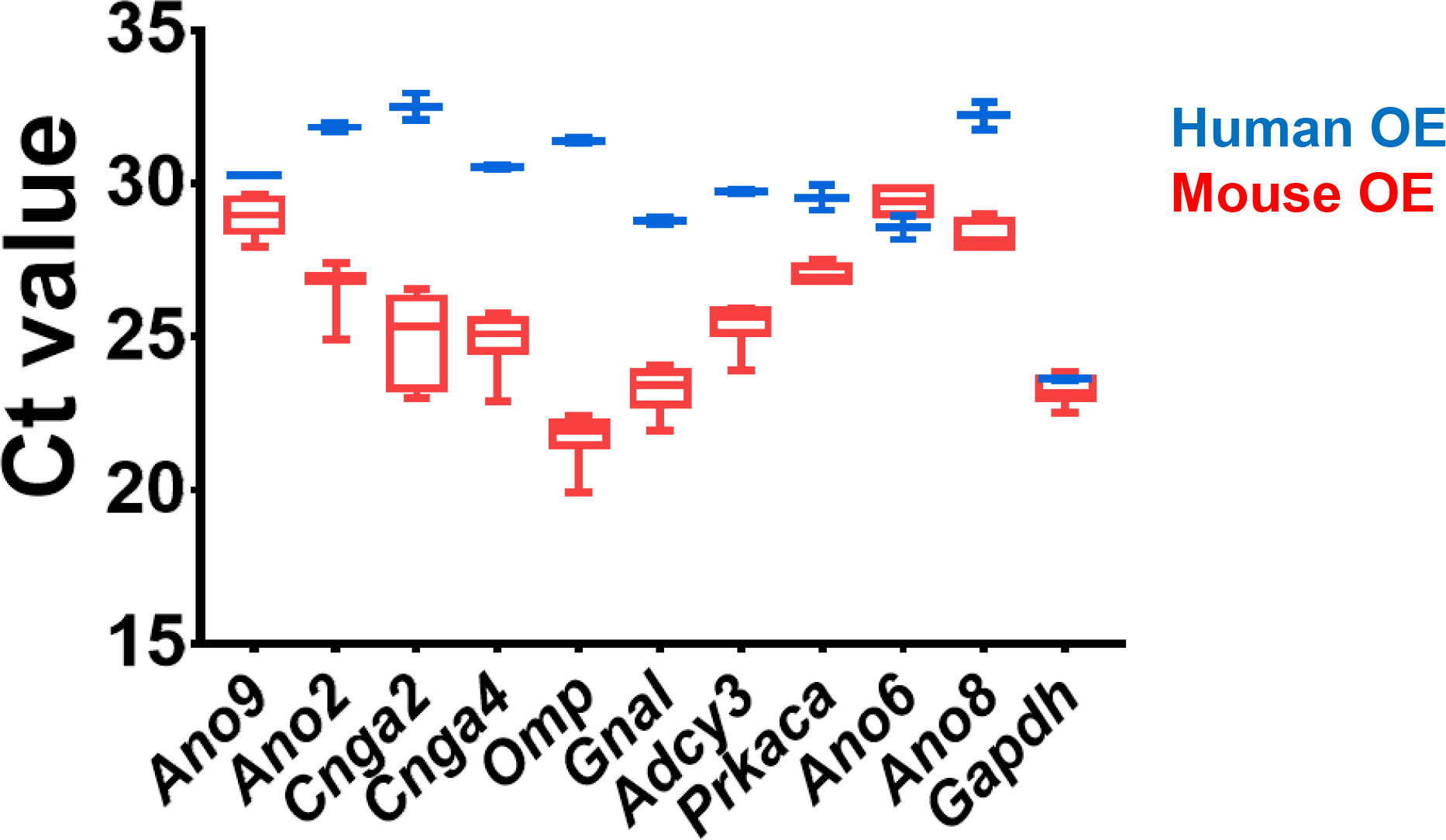
The transcript levels of the Anoctamins and olfactory signal genes in the mouse and human OE. Box plots represent Ct values of various genes in the mouse (red) and human OE (blue) analyzed by quantitative RT-PCR. Global Ct values of the genes tested are shown as the 25th and 75th quartiles (boxes), median (central horizontal line), and minimal/maximal value (vertical bar). Quantitative PCR was performed using cDNAs quantitatively diluted to 100 ng of each sample.

**Extended Data Fig. 2.**
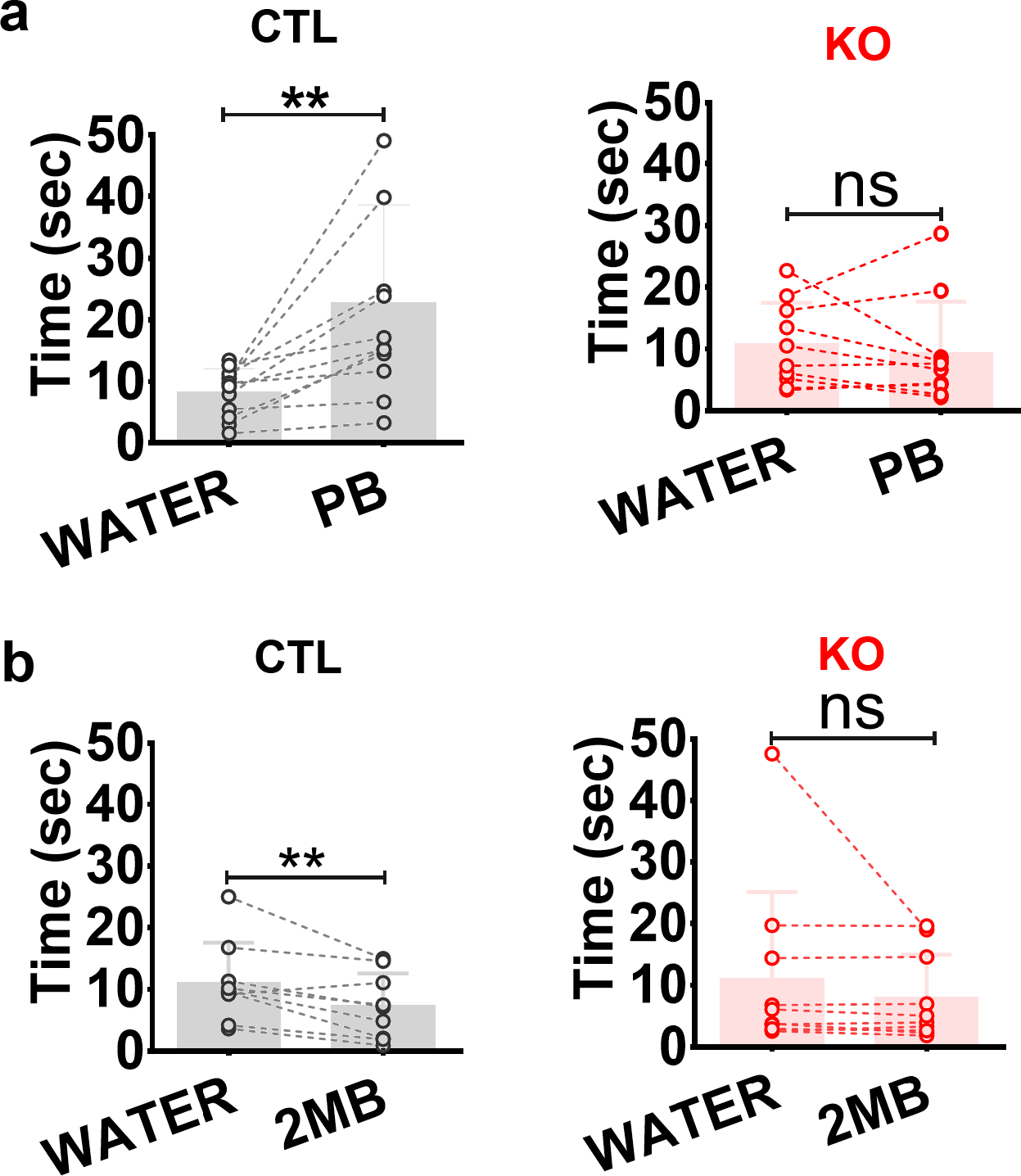
Odorant preference test. Time spent exploring a dish with water and peanut butter (PB) **(a)** or a water and 2-methylbutyric acid (2MB) (**b**) in a 3-minute test. Connected circles represent the same mouse in the test. ** p < 0.01, ns: not significant, Student T-test.

**Extended Data Fig. 3.**
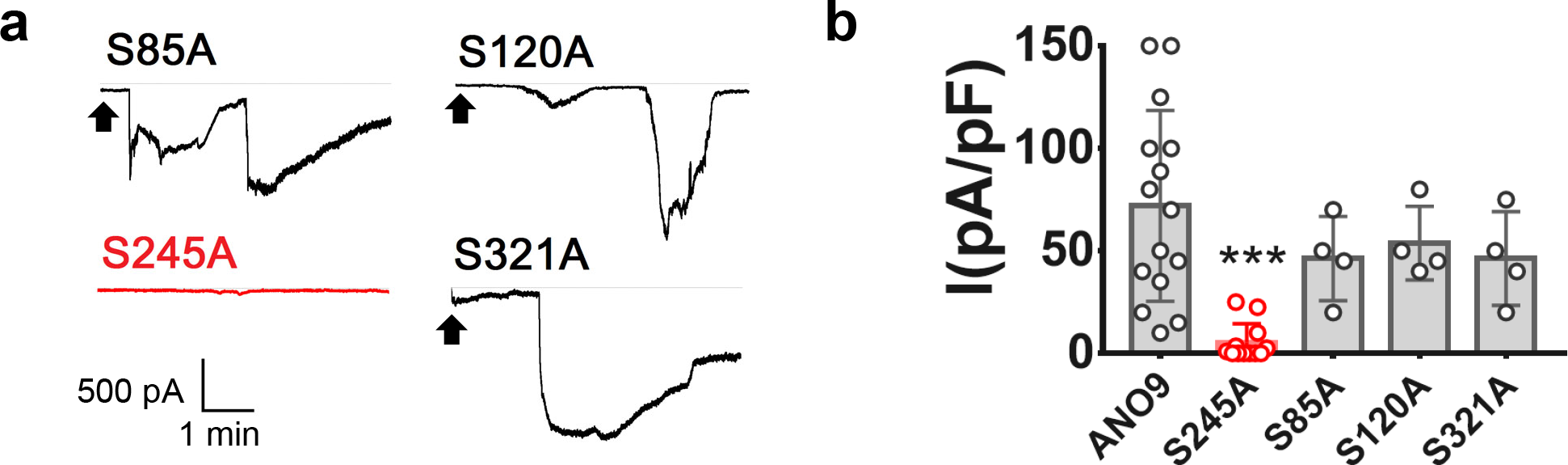
The ANO9 S245A mutant fails to be activated by cAMP. Representative **(a)** and summary **(b)** of whole-cell currents of HEK 293T cells transfected with *Ano9* and each of four mutants (S85A, S120A, S245A, and S321A). Intracellular cAMP evoked robust inward currents in the S85A, S120A or S321A mutant-transfected HEK cells but not in the S245A/HEK cells. *** p < 0.001 compared to control ANO9, One-way ANOVA, Tukey’s post-hoc test.

**Extended Data Fig. 4.**
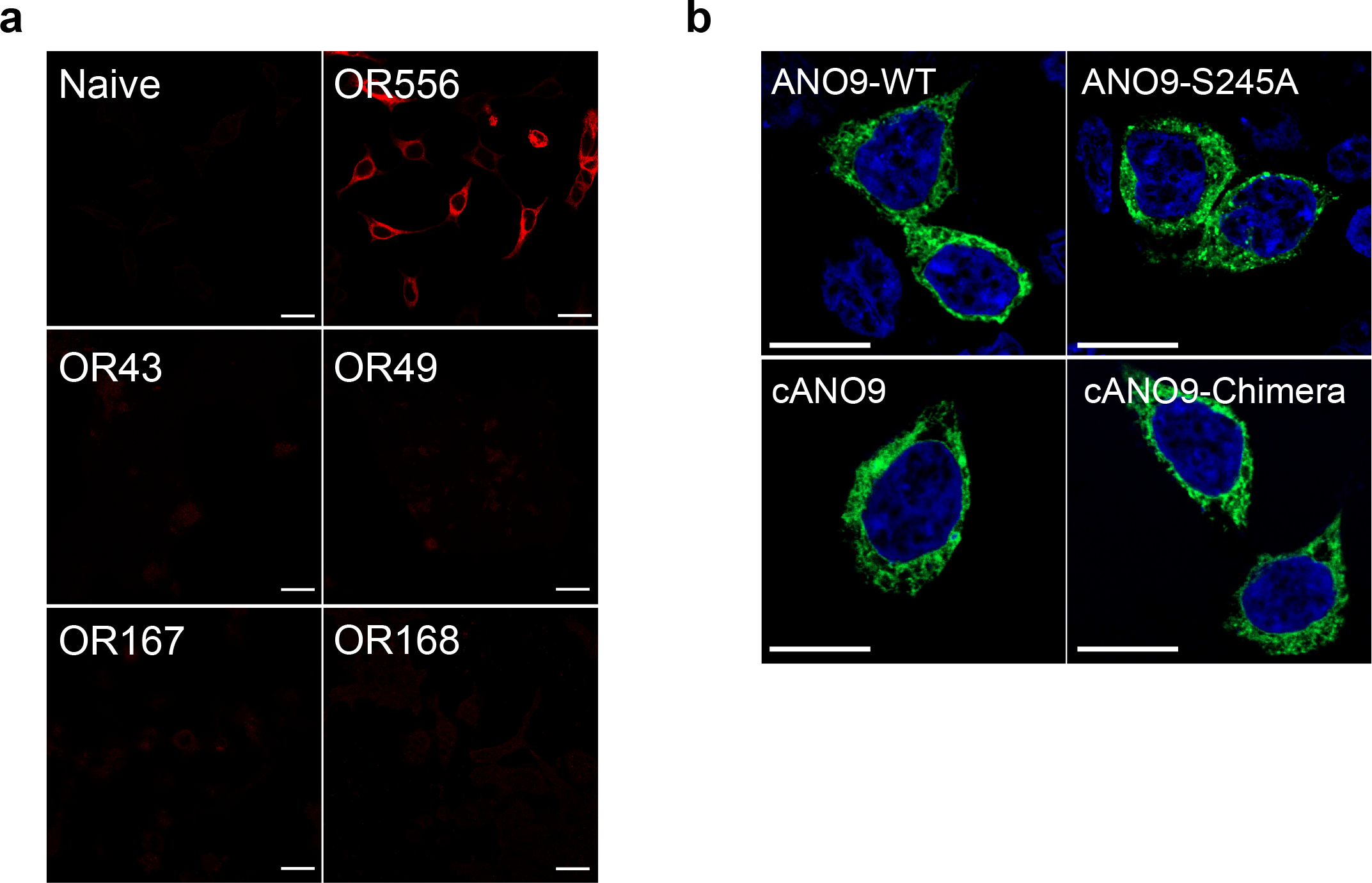
Heterologous expressions of various ORs and *Ano9* mutants in HEK 293T cells. **a,** The expressions of OR556, OR43, OR49, OR167 and OR168 tagged with Myc were checked with Myc antibody in HEK 293T cells. Only OR556 was identified for its expression in the HEK 293T cells. Bars represent 20 μm **b,** GFP-tagged mouse ANO9, S245A, chick ANO9, and chick-mouse ANO9 chimera were transfected to HEK 293T cells.

**Extended Data Fig. 5.**
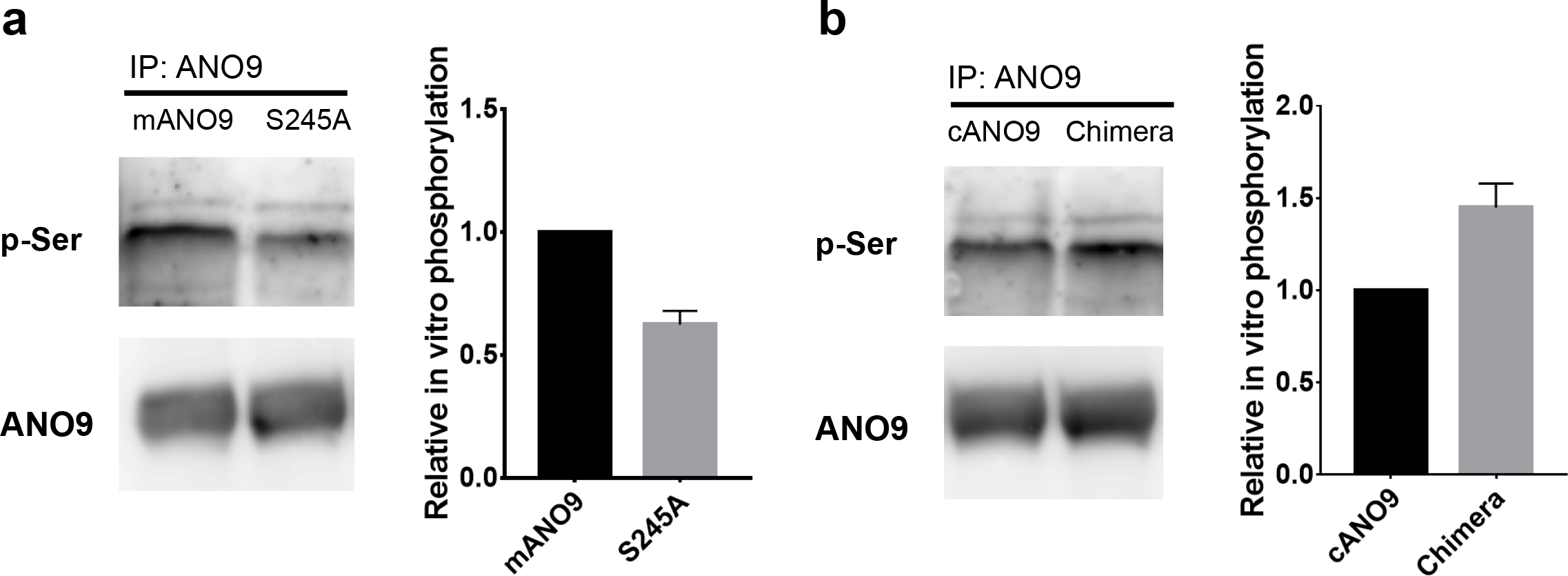
The Ser245 residue is the main phosphorylation site by PKA. Immunoblots by phosphoserine antibody of the precipitants with ANO9 antibody of PKA-phosphorylated ANO9/HEK or mAno9/S245A/HEK cells **(a)** and cAno9/HEK cells **(b)**. The lysates of HEK 293T cells transfected with *Ano9*, the S245A mutant, or chick *Ano9* were precipitated with anti-ANO9 antibodies. The lysates were then phosphorylated after incubation with a purified recombinant catalytic subunit of PKA (2,500 Unit/ml) together with 200 μM ATP. These immunoprecipitants were blotted with anti-phosphoserine antibodies.

**Extended Data Fig. 6.**
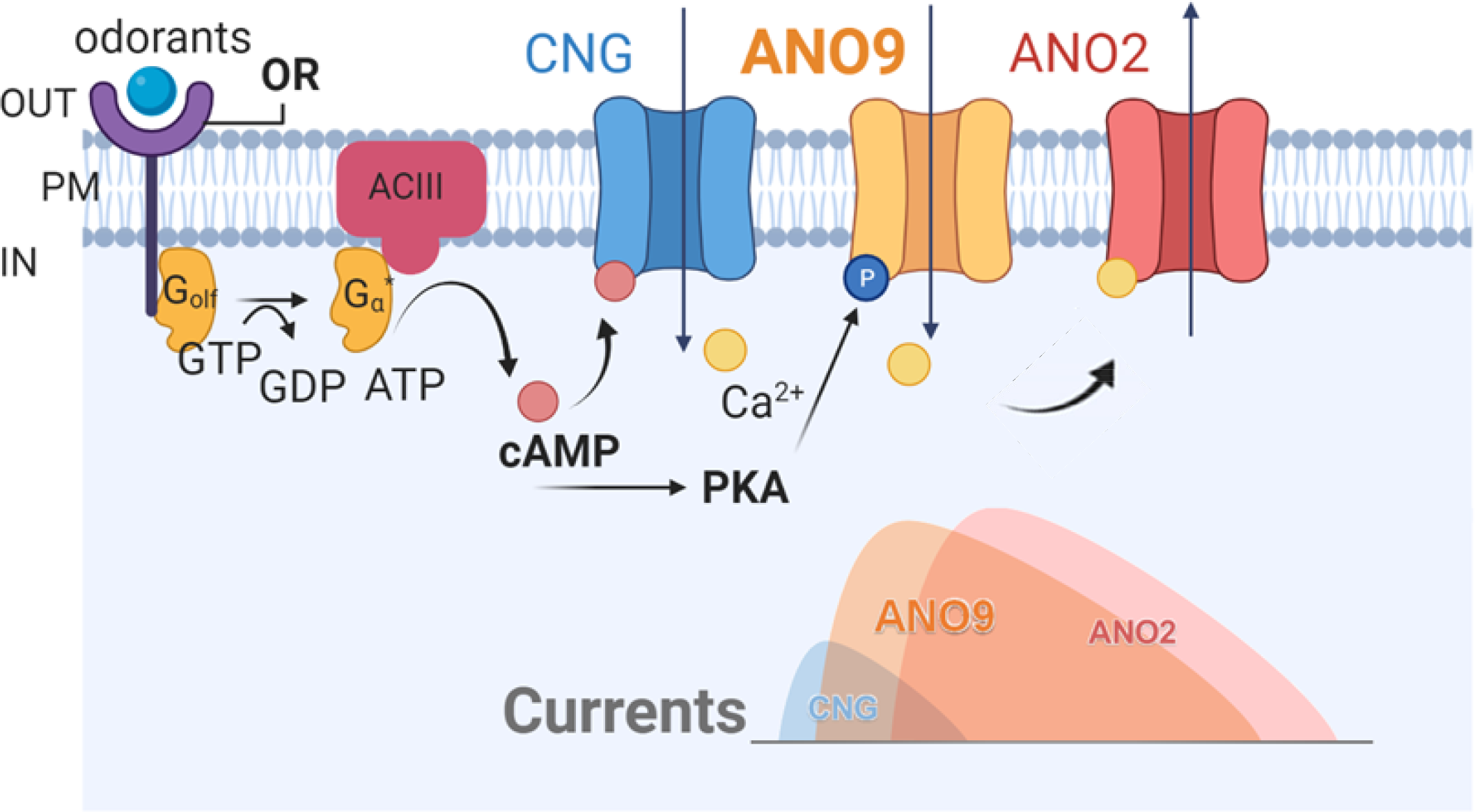
Proposed olfactory signal transduction pathway.

